# GROQ-seq Datasets Across Transcription Factors (LacI, RamR, VanR), T7 RNA Polymerase and TEV Protease

**DOI:** 10.64898/2026.04.15.718744

**Authors:** Aviv Spinner, Shwetha Sreenivasan, James R. McLellan, Svetlana Ikonomova, Dana Cortade, Simon d’Oelsnitz, Kristen Sheldon, Olga Vasilyeva, Nina Y Alperovich, Anjali Chadha, Lily Nematollahi, Andi Dhroso, Zach Sisson, Douglas Densmore, Corey M. Hudson, Erika DeBenedictis, Peter J. Kelly, Amanda Reider Apel, David Ross, Catherine Baranowski

## Abstract

Predicting any protein’s function from its sequence alone would be a significant breakthrough in molecular biology. Although machine learning approaches have sought to tackle this, their limited generalizability reflects the absence of sufficiently large, open, diverse, and unified datasets. To address this data gap, we developed a high-throughput experimental platform called GROQ-seq (**Gro**wth-based **Q**uantitative **Seq**uencing). In GROQ-seq, a protein’s function can be linked to a sequencing-based readout that enables scalable characterization of large variant libraries in *Escherichia coli*. Here, we present pilot datasets demonstrating its performance across three distinct protein function classes: transcription factors, polymerases, and proteases. The objective of this report is to present the datasets and to provide users with a clear and transparent characterization of their properties, including both the strengths and limitations.

## Introduction

GROQ-seq is a pooled *E. coli*, growth-based assay that links protein function to cellular fitness by coupling variant activity to antibiotic resistance. This is achieved using a barcoded plasmid-based genetic circuit expressing the gene of interest, where its function modulates the activity or expression of dihydrofolate reductase (DHFR) (Supplemental Figures 3A, 4A, 5A). Since DHFR confers resistance to trimethoprim (TMP), differences in protein function lead to corresponding differences in TMP resistance. Each variant is uniquely barcoded, allowing next-generation sequencing to quantify barcode abundance, which serves as a proxy for variant growth (fitness) in the presence of TMP.

Before conducting a GROQ-seq experiment, the protein function of interest must first be onboarded. This process begins with plasmid design, where a gene circuit is constructed to link protein function to growth. The circuit is then validated for responsiveness and optimized to expand its dynamic range. Next, the system is calibrated using barcoded variants that span the full range of functional activity and are measured individually before being incorporated into pooled assays. The pooled assay is ideally validated by confirming strong agreement between singleplex and pooled measurements of these calibration variants. (See the Transcription factor^2^, T7 RNA Polymerase^3^ and TEV Protease^4^ GROQ-seq Technical Bulletins for more details).

A key feature of GROQ-seq is the inclusion of a barcoded calibration ladder in every pooled experiment. This enables quantitative, reproducible measurements across different experimental batches as well as the conversion of relative fitness into quantitative protein function values (See Supplemental Figure 2 for an overview of the data analysis pipeline). For additional details on the assay, see Spinner et al ^1^.

To demonstrate its broad applicability, the first three protein functions tested by GROQ-seq were transcription factors (TF), RNA polymerase (RNAP) and proteases. These represent three progressive layers of information processing in the cell: regulatory sensing, transcription, and post-translational processing. Together, these diverse functional classes provide a robust foundation of inputs to illustrate the versatility and generalizability of the GROQ-seq pipeline.

### GROQ-seq Data on Transcription Factors, TEV Protease and T7 RNA Polymerase

#### Transcription Factors

In this report, we present results from GROQ-seq experiments characterizing the function of three bacterial TFs: LacI, RamR, and VanR. In GROQ-seq, the functional readout depends on the protein class and for TFs this corresponds to regulation of gene expression. Each TF binds its cognate DNA operator, controlling transcription of a DHFR gene (Supplemental Figure 3A, for details, please refer to the Technical Bulletin for GROQ-seq Transcription Factor Function Assay^2^).

To quantify each TF’s response to its corresponding small-molecule ligand, we used the GROQ-seq assay to measure two functional states: the uninduced transcription rate (in the absence of ligand) and the induced transcription rate (in the presence of a high ligand concentration) (Supplemental Figure 3B). Results for both the uninduced and induced transcription rates are included in the data release for the TFs^5–7^. To provide an initial overview of the data, we focus on results for uninduced transcription, the log_g0 column from the released data tables, which is the base-10 logarithm of the transcription rate in the absence of ligand. Uninduced transcription is expected to be inversely related to the strength of the interaction between the transcription factor and its cognate operator^8–12^. So, lower log_g0 values correspond to higher “activity” (stronger binding and therefore repression). Across variants, transcription rates measured in the GROQ-seq assay span approximately 2.5 orders of magnitude, and the calibration ladder variants were selected to cover this full dynamic range. For each TF, we investigated substitution, deletion, and insertion libraries (SSVL), site saturation mutagenesis libraries (SSM), and error-prone PCR multimutant libraries (epPCR), achieving broad mutational coverage.

#### Transcription Factor Variant Libraries

The GROQ-seq data release includes sequence-function data for 44,914 LacI variants, 32,593 RamR variants, and 23,966 VanR variants. For SSVL libraries, we observed 57.3% of possible single-mutant variants for LacI, 80% for RamR, and 48.6% for VanR (Table 1). Total variant counts reported here reflect deduplicated variants across library types, whereas counts in Table 1 are reported per library and therefore will not sum to the deduplicated total due to overlap between libraries.

**Table 1:** Transcription factor library statistics.

| Protein | Library Type | Total Unique Variants | Total Variants with Single Mutation | Total Possible Designs | Percent of total designs tested* | Mean Number of Barcodes Per Variant | Median Number of Mutations per Variant | Mean Number of Mutations per Variant |
| --- | --- | --- | --- | --- | --- | --- | --- | --- |
| LacI | epPCR | 36,656 | 1,624 | - | - | 1.3 | 3 | 3.1 |
|  | SSM | **54** | 19 | **8,000** | 0.7 | 6.9 | 2 | 1.8 |
|  | SSVL | 9,769 | **8,276** | **14,440** | 57.3 | 3.9 | 1 | 1.2 |
| RamR | epPCR | 25,157 | 999 | - | - | 1.5 | 3 | 2.8 |
|  | SSM | **1,181** | 24 | **8,000** | 14.8 | 1.2 | 3 | 2.8 |
|  | SSVL | 7,247 | **6,239** | **7,800** | 80 | 4.8 | 1 | 1.1 |
| VanR | epPCR | 18,639 | 915 | - | - | 1.8 | 2 | 2.6 |
|  | SSM | **13** | 2 | **8,000** | 0.2 | 1.5 | 3 | 2.3 |
|  | SSVL | 6322 | **4,765** | **9,800** | 48.6 | 8.1 | 1 | 1.3 |

\*Percentage of designs tested is calculated as the number of variants observed from that library divided by the total possible library space (bolded values).

Some positions were missing from the gene library synthesis used for the SSVL libraries. For RamR those missing positions are T44, L45 and F46, as well as Y177, I178, A179, and L180. For VanR those missing positions are A233 and P234.

The GROQ-seq plasmid circuit for TFs showed mild toxicity at high transcriptional output, causing variants with high uninduced transcription to drop out of the dataset. This bias is strongest in VanR and weakest in RamR. We are re-optimizing the GROQ-seq measurement to reduce this bias in future datasets.

Despite this, the TF GROQ-seq experiment has been shown to produce reproducible, quantitative sequence–function measurements across barcodes, runs, and independent facilities^1^.

#### Transcription Factor GROQ-seq Functional Data

For single-mutant TF variants, GROQ-seq results for uninduced transcription are shown in Figures 1-4 for substitutions and deletions and Figures 5-7 for insertion variants. Within Figures 1-3, heatmaps summarize position-specific functional effects (Figures 1A-3A), and the average effect of substitutions are mapped onto protein structures to visualize spatial patterns (Figures 1B-3B). In Figure 4, histograms show the distribution of single-mutant function values, and donut plots capture the proportion of functional categories for each TF.

**Figure 1:**
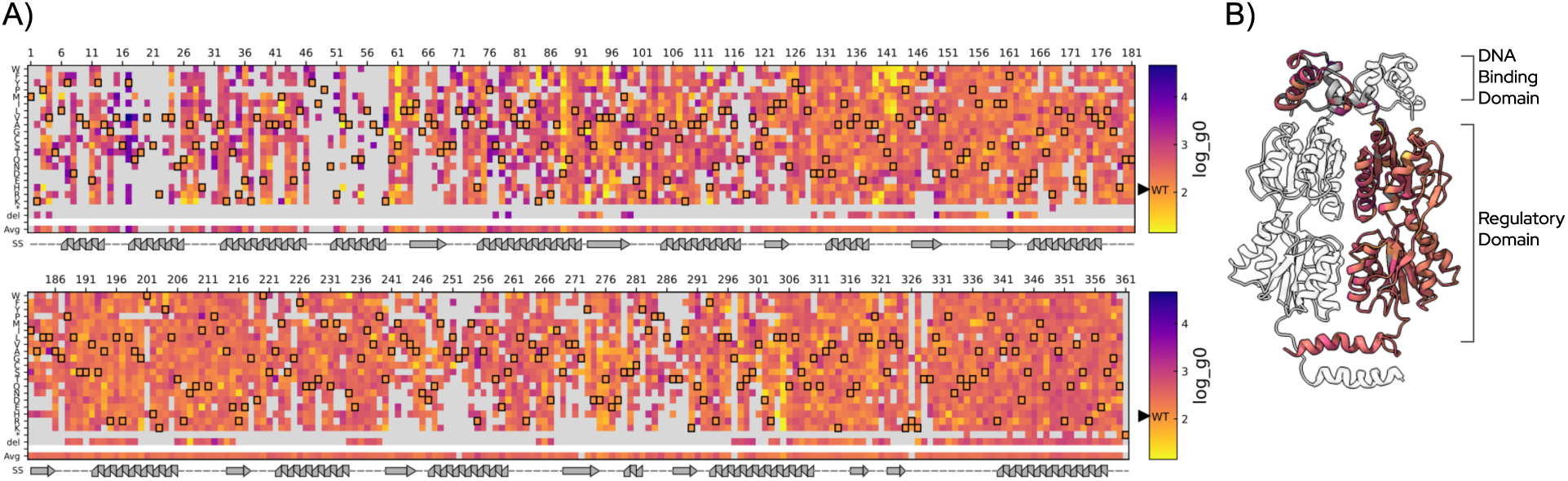
LacI GROQ-seq data for substitution and deletion variants.

**Figure 2:**
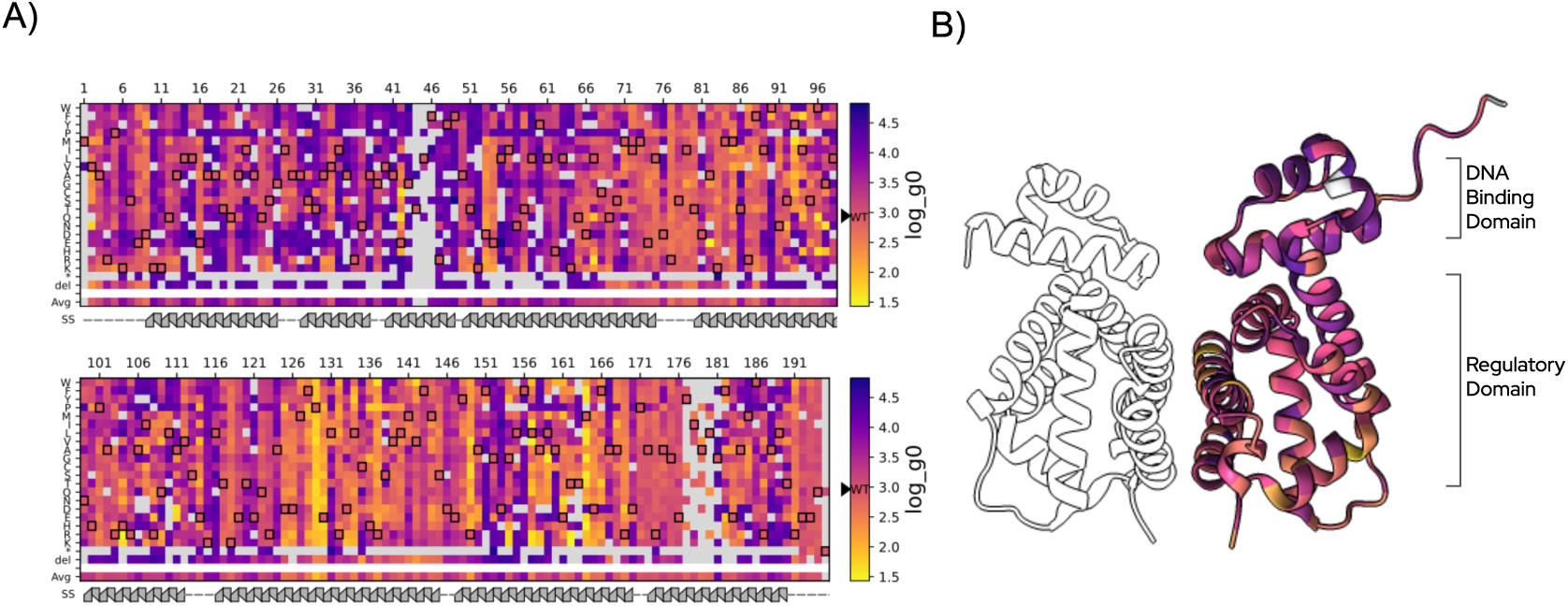
RamR GROQ-seq data for substitution and deletion variants.

**Figure 3:**
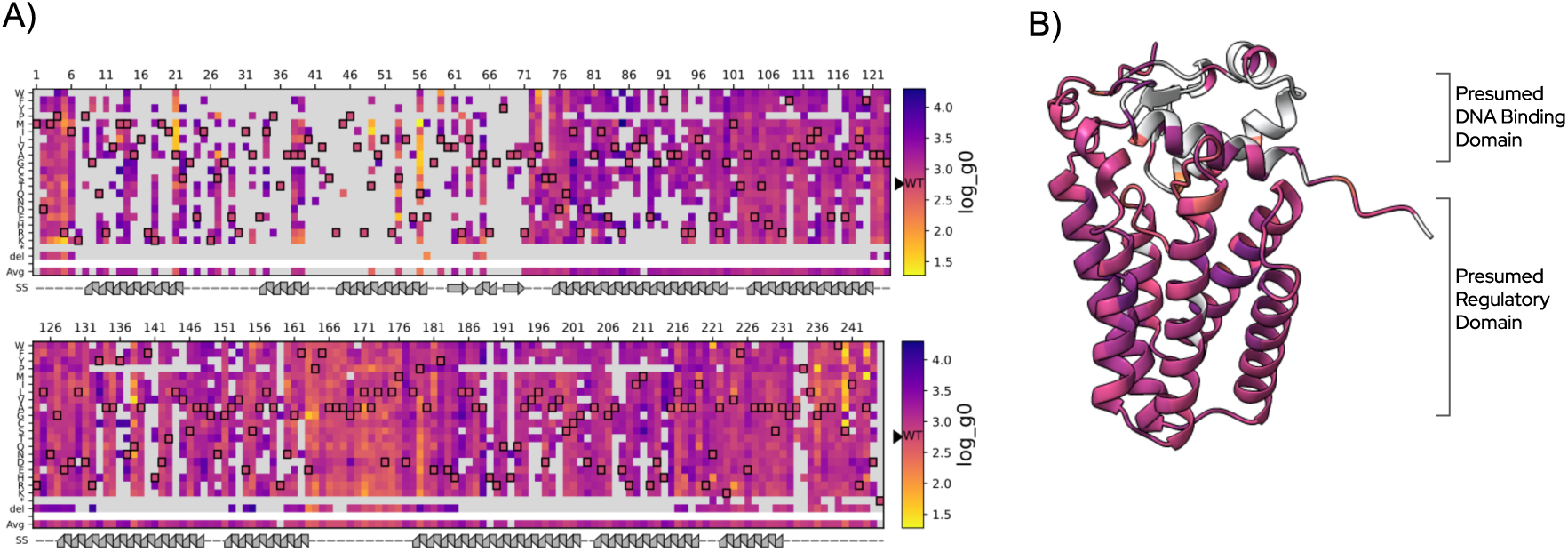
VanR GROQ-seq data for substitution and deletion variants. Figures 1-3: TF GROQ-seq data for substitution and deletion variants (**1A, 2A, 3A**) Heatmaps showing the function values for the base-10 logarithm of the uninduced transcription (log_g0) for single-mutant variants of LacI, RamR and VanR. Uninduced transcription is expected to be inversely related to the binding strength of the TF to its cognate operator, so lower log_g0 values correspond to higher “activity” (stronger binding). Each heatmap is split into two panels stacked vertically, with each subpanel spanning half the total length of the protein. The x-axis represents residue position, and the y-axis has the amino acids grouped by physicochemical properties of their side chains followed by the stop codon (*), deletion (del), and the average effect of mutations at each position (avg). Each cell is colored on a continuous scale spanning the observed range of function values shown on the right with the value of the wild-type marked by the filled black triangle labeled “WT”. The wild-type residue at each position is marked with a black outline in the heatmap. Missing data (e.g., missing variants from the library synthesis), and the average cell for positions with less than five reported substitutions are rendered in gray. A secondary structure annotation (helix, strand, loop) is drawn below each panel. (**1B, 2B, 3B**) Protein structure images with residues colored by the average log_g0 value for single amino acid substitutions (LacI PDB: 1LBG^13^; RamR PDB: 3VVX^14^; VanR AlphaFold predicted structure AF-Q9A5Q5-F1^15^). In each case, one monomer of the biological assembly is colored by the per-residue average functional value (with positions containing <5 mutations rendered in white) while the remaining subunits are rendered transparent to provide structural context. For LacI and RamR, the DNA-binding domain and the regulatory domains are marked to the right. No experimental structure exists for VanR, so the image shows a single monomer from the AlphaFold predicted structure with the presumed domains marked to the right. For each structure, the DNA-binding domain is oriented toward the top of the structure.

**Figure 4:**
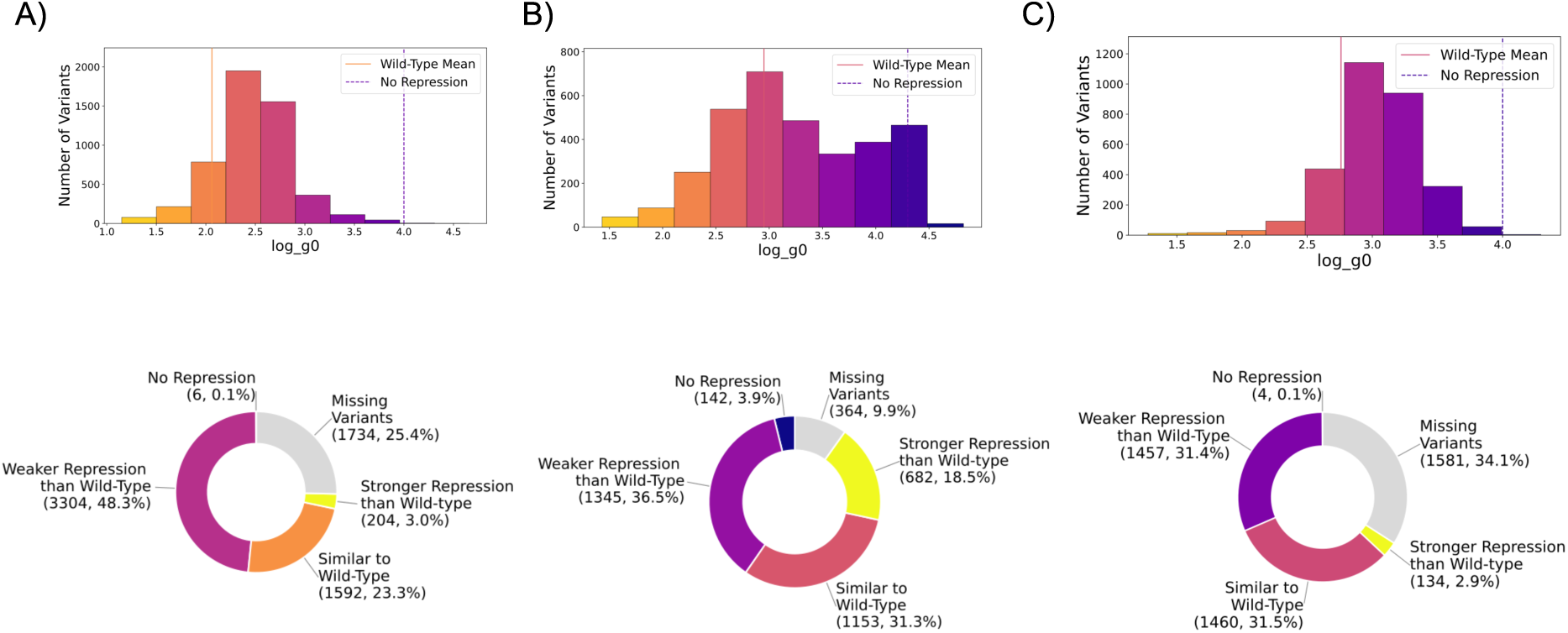
Comparison of TF single substitution variants to wild-type function Histograms and donut plots of log_g0 values for single substitution variants of LacI, RamR and VanR. (**A, B, C top**) The histogram plots show the range of the measured outcomes split into 10 equal-sized intervals on the x-axis and the number of variants sampling each interval on the y-axis. Each bar is colored using the same colormap as the corresponding heatmaps in Figures 1-3. The wild-type mean is indicated by the solid line; the threshold for the no-repression category (dashed line) is set to the mean value of the induced wild-type. The colors of both the lines correspond to their values in the respective heatmaps in Figures 1-3; (**A, B, C bottom**) The donut plots shows the proportion of single substitutions with log_g0 values categorized relative to the wild-type: No repression (high log_g0), weaker repression than wild-type (log_g0 values between no-repression threshold and similar-to-wild-type), similar to wild-type (within 2-fold of wild-type mean on either direction), stronger than wild-type (low log_g0). Missing variants are shown in gray. Because of the toxicity associated with high log_g0, most missing variants are expected to be in the no-repression category.

**Figure 5:**
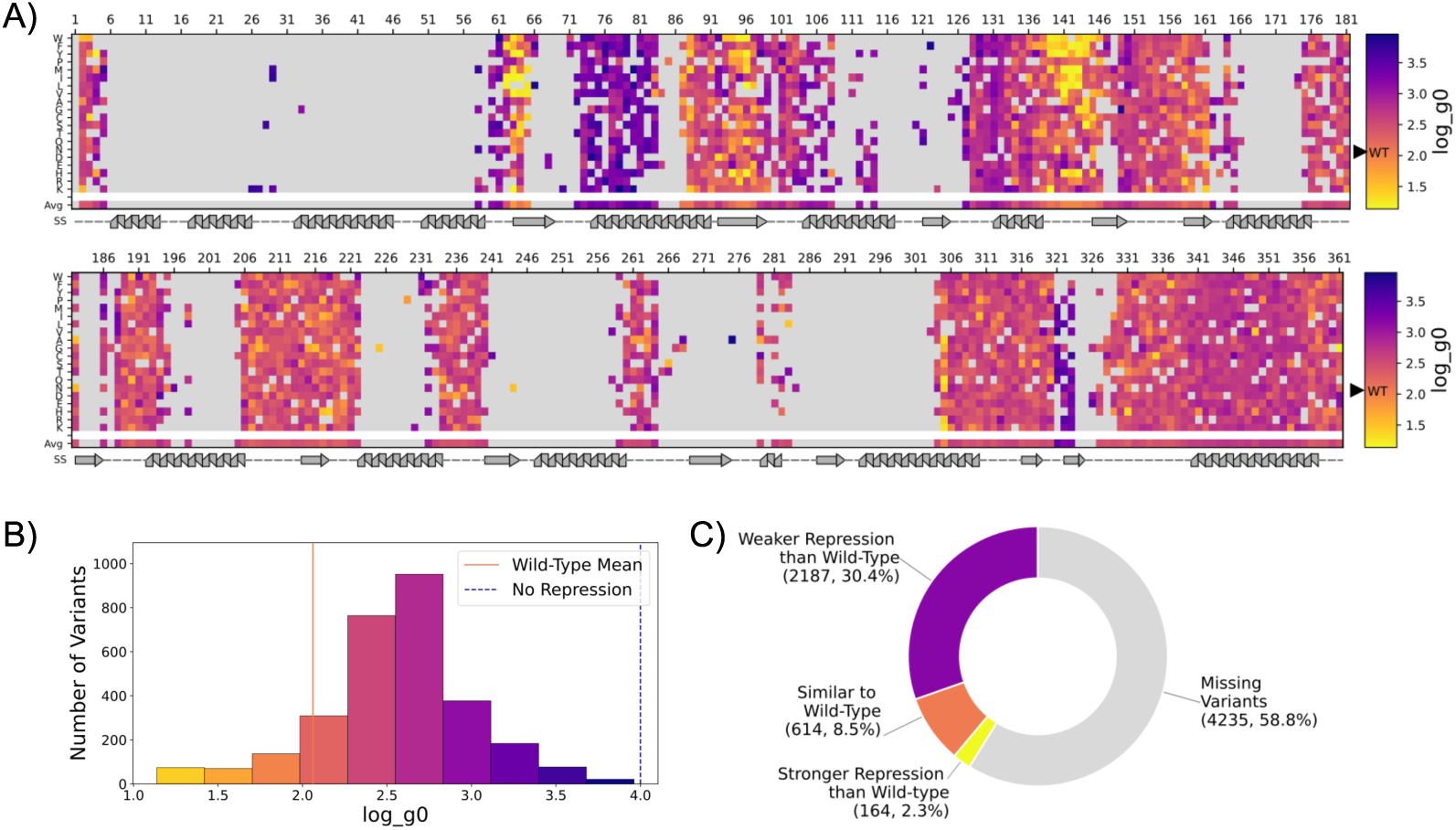
GROQ-seq data for LacI insertion variants.

Across all three TFs, a surprising fraction of single substitutions result in activity (repression) similar to or stronger than the wild-type: 26.3% for LacI, 49.8% for RamR, and 34.4% for VanR (where similar to wild-type is defined as within 2-fold of the wild-type repression, i.e., within ± log_10_(2) of the wild-type value for log_g0) (Figure 4). Interestingly, these results align with the conclusion of von der Dunk et al. that, for naturally occurring proteins, about 25% of single substitutions result in “… an identical backbone structure as the wildtype…”^16^. The trend across the different TFs also matches the result from von der Dunk et al., that proteins with a higher fraction of alpha-helical secondary structure tend to be more robust to perturbation from mutations: The percent alpha-helical structure is 46%, 84%, and 66% for LacI, RamR, and VanR, respectively. This ordering of mutational tolerance (RamR > VanR > LacI) mirrors the increasing fraction of α-helical content in these proteins, consistent with the idea that higher α-helical structure contributes to greater robustness to single substitutions.

In each TF, there are notable regions where mutations tend to increase the repression strength (i.e., lower log_g0 than wild-type, stronger interaction with the DNA operator). Some of these regions are near the DNA binding domain (top of structures in Figures 1B-3B), but not in direct contact with the DNA operator, indicating a mechanism that is likely related to allosteric communication between protein domains. Examples include positions S61 in LacI (in a loop that is part of the hinge between the DNA-binding and regulatory domains), N142 in LacI (in a loop of the regulatory domain that faces toward the DNA), and S240 in VanR (in the C-terminal loop that is folded back toward the DNA). At each of these positions, mutations to most hydrophobic residues result in stronger repression (yellow color in Figure 1A, 3A). More strikingly, in RamR, there are several positions in the middle of alpha helices and on the opposite side of the protein from the DNA-binding domain where a broad set of mutations increase the repression strength (e.g., P129, E130, R137, M164, in Figure 2A).

Across all three TFs, insertions recapitulate the patterns seen for substitutions, with the same regions and physicochemical biases driving increased repression. For example, in LacI, many hydrophobic insertions between positions S61 and G65 or between A138 and P144 increase repression (Figure 5A). Similarly, in VanR hydrophobic insertions between G236 and A237 increase repression (Figure 7A). And, in RamR, a broad set of insertions between L131 and R137 or between A167 and P171 increase repression (Figure 6A).

**Figure 6:**
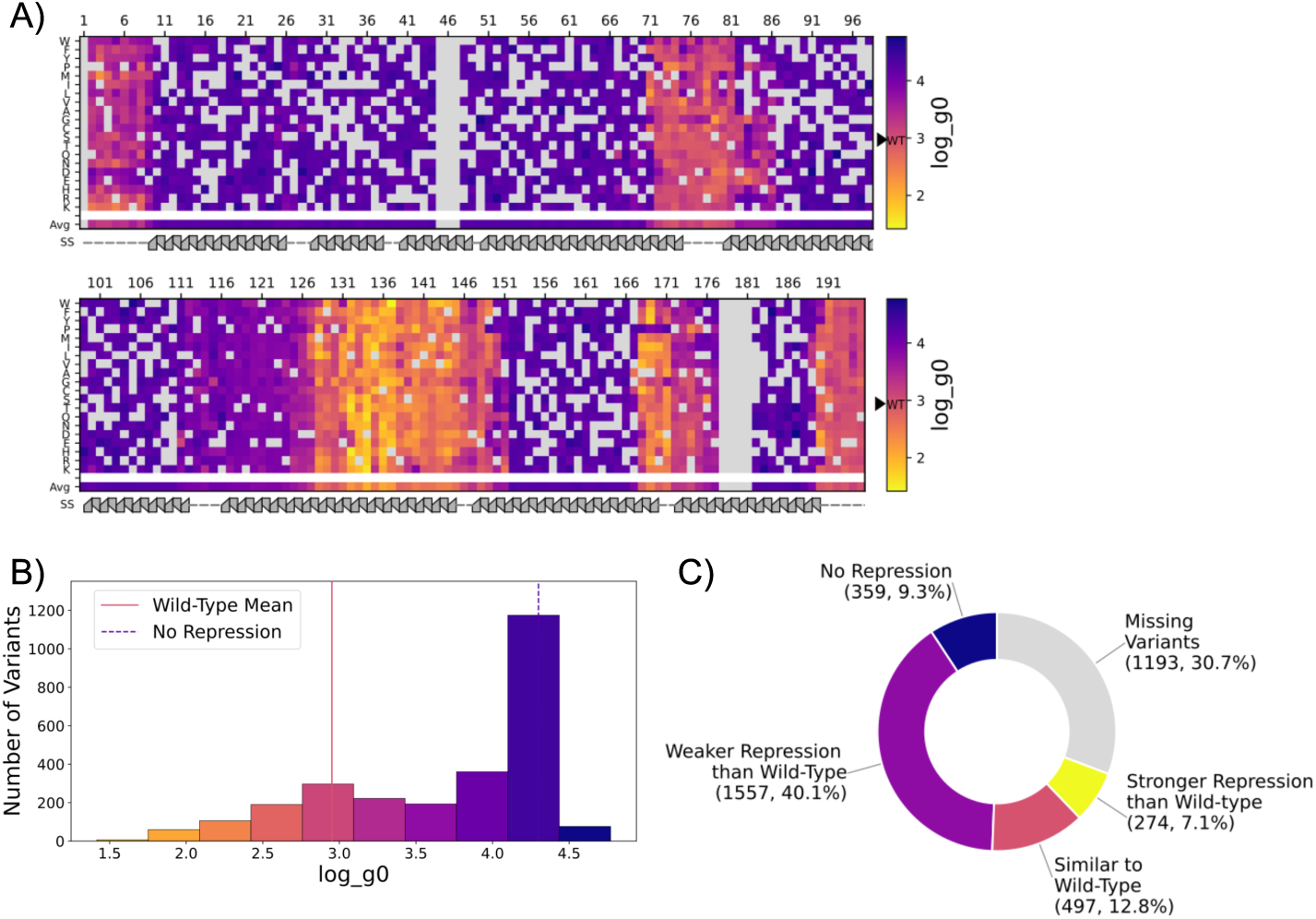
GROQ-seq data for RamR insertion variants.

**Figure 7:**
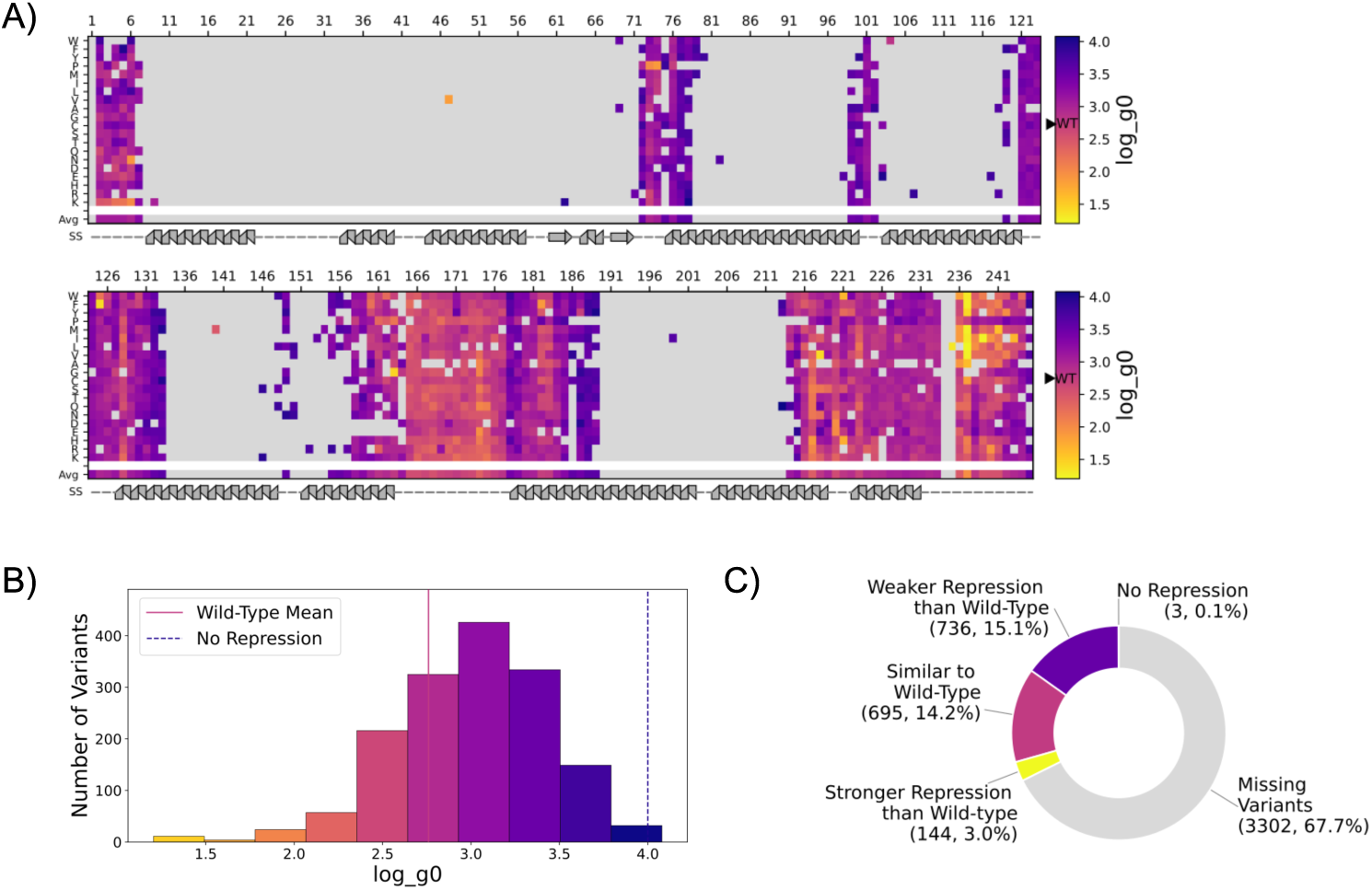
GROQ-seq data for VanR insertion variants. Figures 5-7: TF GROQ-seq data for insertion variants (**5A, 6A, 7A**) Heatmaps showing the function values for the base-10 logarithm of the uninduced transcription (log_g0) for single-insertion variants of LacI, RamR and VanR. Uninduced transcription is expected to be inversely related to the binding strength of the TF to its cognate operator, so lower log_g0 values correspond to higher “activity” (stronger binding). Each heatmap is split into two panels stacked vertically, with each subpanel spanning half the total length of the protein. The x-axis represents residue position, and the y-axis has the amino acids grouped by physicochemical properties of their side chains followed by the average effect of insertions at each position (avg). Each cell is colored on a continuous scale spanning the observed range of function values shown on the right. Missing data (e.g., missing variants from the library synthesis), and the average cell for positions with less than five reported insertions are rendered in gray. A secondary structure annotation (helix, strand, loop) is drawn below each panel. (**5B, 6B, 7B**) Histograms showing the range of the measured outcomes for insertions for LacI, RamR and VanR. The histogram plots show the range of the measured outcomes split into 10 equal-sized intervals on the x-axis and the number of variants sampling each interval on the y-axis. Each bar is colored using the same colormap as the heatmaps in the respective panel A’s. The wild-type mean is indicated by the solid line; the threshold for the no-repression category (dashed line) was set to the mean value of the induced wild-type. The colors of both the lines correspond to their values in the heatmaps in the respective panel A’s. (**5C, 6C, 7C**): The donut plots shows the proportion of single insertions with log_g0 values categorized relative to the wild-type for LacI, RamR, and VanR: No repression (high log_g0), weaker repression than wild-type (log_g0 values between no-repression threshold and similar-to-wild-type), similar to wild-type (within 2-fold in either direction from the wild-type mean), stronger than wild-type (low log_g0). Missing variants are shown as gray in the donut plots. Because of the toxicity associated with high log_g0, most missing variants are expected to be in the no-repression category.

To assess combinatorial effects beyond single mutations, epPCR libraries that contain multi-mutant variants were assayed via GROQ-seq. Most TF epPCR variants contained two mutations, with a longer tail extending to higher mutation counts (Figure 8). The average number of mutations per variant was 3.1 for LacI, 2.8 for RamR, and 2.6 for VanR. Consistent with expectations for random mutagenesis, TF repression generally decreases (log_g0 increases) as the number of mutations increases, reflecting the cumulative destabilizing or disruptive effects of multiple amino acid substitutions (Figure 8B, D, F). Notably, a substantial fraction of variants with up to five or six mutations remained functional. This likely arises from both mutational robustness and selection against highly expressed, non-functional variants due to circuit toxicity.

**Figure 8:**
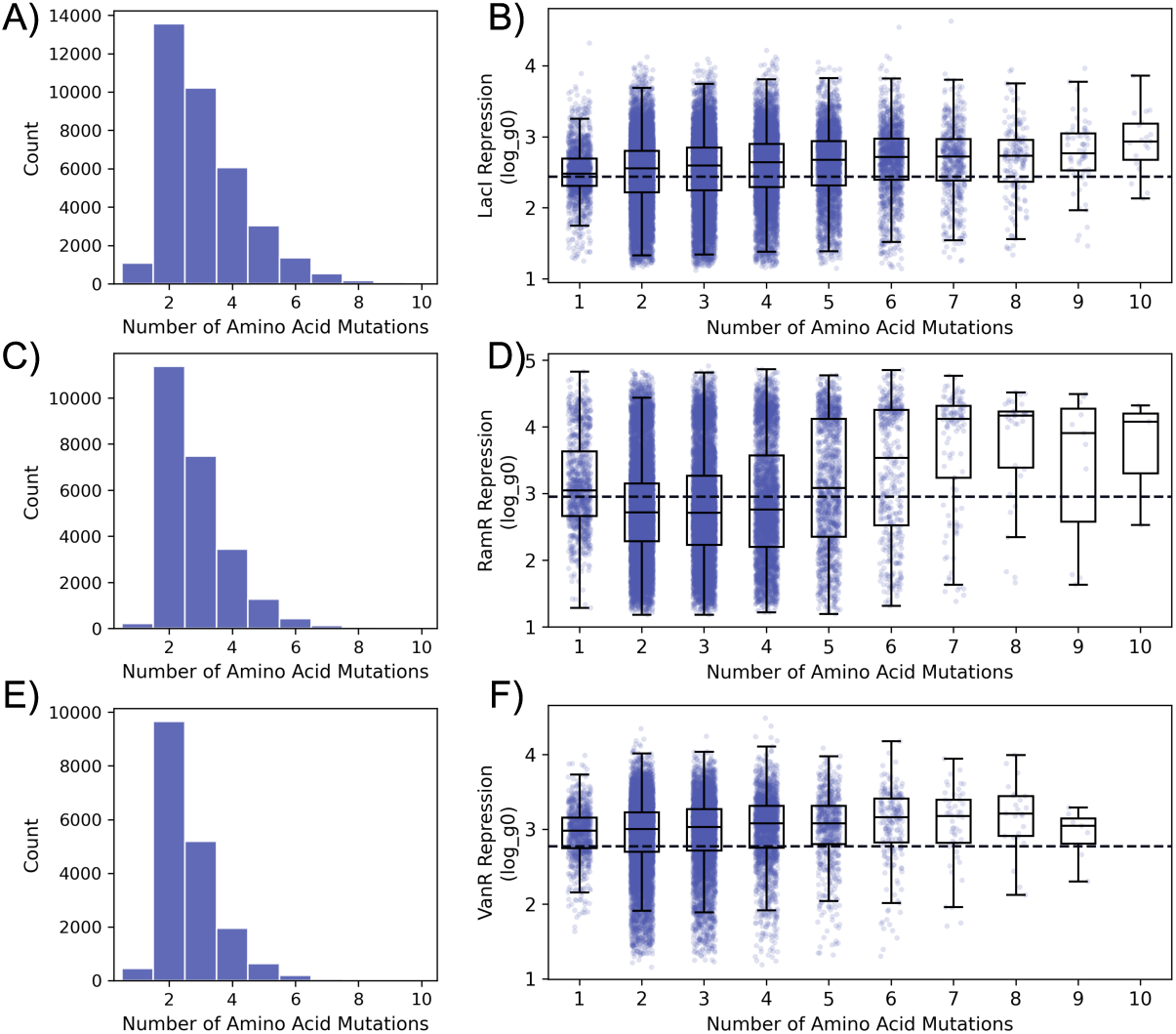
TF epPCR library GROQ-seq data. (**A, C, E**) Distribution of number of variants (LacI, RamR, VanR); (**B, D, F**) Repression levels for epPCR variants, dashed-line represents wild-type uninduced transcription activity (LacI, RamR, VanR), whiskers indicate ±1.5× inter-quartile range.

#### Comparison to Existing Datasets

To highlight where GROQ-seq builds upon available data, we compared the GROQ-seq TF datasets to existing mutagenesis data. While comparisons to existing datasets show areas of agreement, observed differences highlight the sensitivity of sequence–function relationships to assay context and emphasize the need for unified, reproducible measurement frameworks.

In one of the earliest comprehensive mutagenesis studies for any protein, Suckow et al., used a semi-quantitative method to classify LacI function into four categories for different levels of transcription, ranging from a score of 1 (similar to wild-type) to 4 (high transcription, i.e., no repression)^17^. Comparison with our GROQ-seq result for log_g0 shows weak correlation (Figure 10A, Spearman ⍴ = 0.27) (Figure 9A).

**Figure 9:**
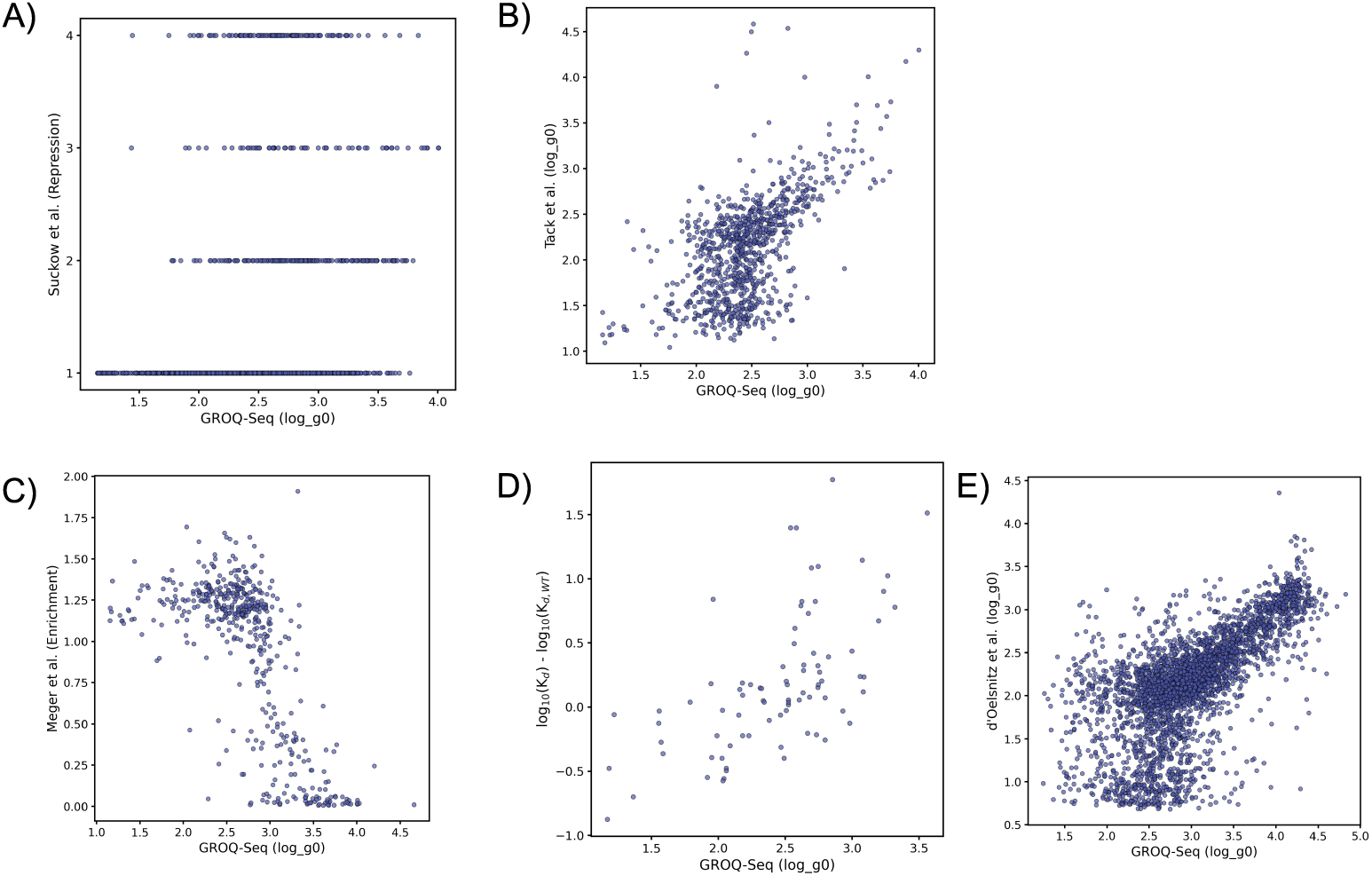
Comparing TF GROQ-seq data with previously published literature data. GROQ-seq data compared to (**A**) Suckow et al., repression data for LacI^17^; (**B**) Tack et al., log_g0 values for LacI^18^; (b) Meger et al., enrichment data for LacI^19^; (**D**) Dissociation constants for LacI from the AlloRep database^20^ (base-10 logarithm of K_d_ for the LacI-operator interaction normalized by the base-10 logarithm of wild-type K_d_); (**E**) d’Oelsnitz et al., log_g0 values for RamR^21^.

**Figure 10:**
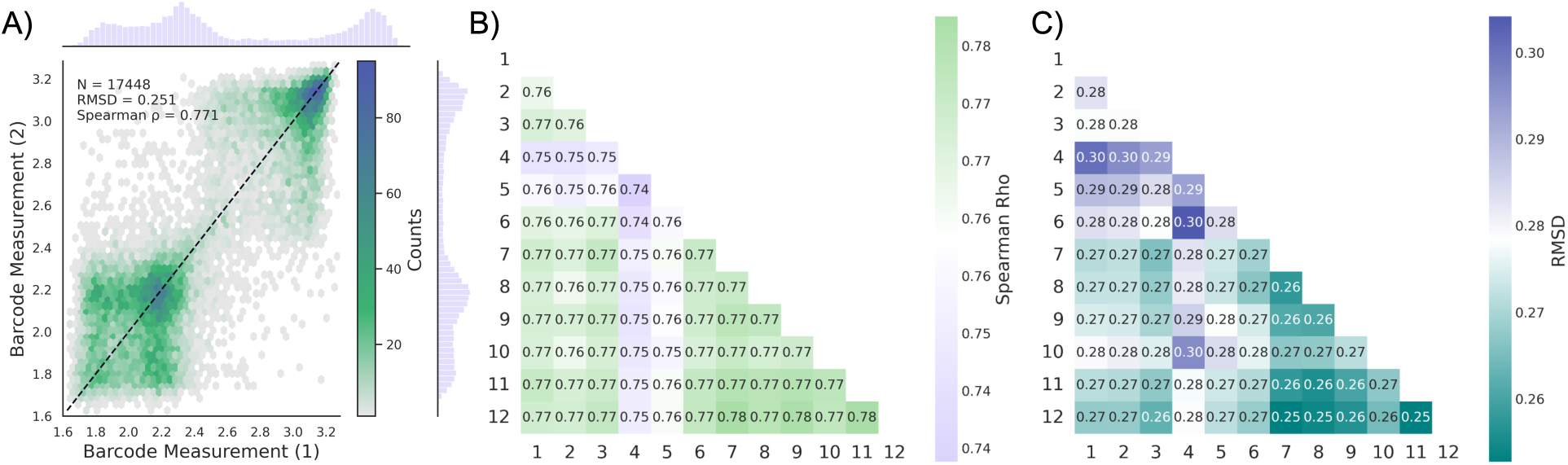
T7 RNAP variant library reproducibility. (**A**) Correlation of data from independent barcodes in a single replicate (replicate 12); (**B, C**) Reproducibility metrics comparing each barcode across 12 identical replicates (Spearman correlation coefficient, Root mean squared deviation [RMSD]).

Tack et al., measured the sequence–function relationships of over 50,000 LacI variants from an epPCR library using an assay that served as a precursor to GROQ-seq^18^. They reported a log_g0 function value (base-10 logarithm of uninduced transcription), enabling more direct comparison with the GROQ-seq dataset reported here. For 899 LacI variants present in both datasets, most of the log_g0 results from Tack et al. were below the dynamic range for the measurement. Nevertheless, comparison with the GROQ-seq result reported here still shows an approximately linear correlation with Spearman ⍴ = 0.54 and Pearson r = 0.59 (Figure 9B).

In another mutagenesis study of LacI, Meger et al. reported growth-based enrichment measurements for 1,122 single amino acid substitutions in the first 60 positions in the N-terminal DNA-binding domain^19^. Comparison with GROQ-seq log_g0 results shows a non-linear correlation for the 483 mutants that overlapped between the two datasets, with Spearman ⍴ = -0.64 (Figure 9C; negative correlation because of differences in assay design). The distribution of data is bimodal for Meger et al. but more continuous for the GROQ-seq result. Also, the relationship between the Meger et al. enrichment scores and the GROQ-seq log_g0 values seems to follow a sharp sigmoidal curve with most of the changes in enrichment occurring over a narrow log_g0 range (between 2.7 and 3.5), indicating a wider dynamic range for the GROQ-seq measurement.

To probe whether GROQ-seq data correlates to existing biophysical data, we used the AlloRep database to obtain *K_d_* values for LacI-operator binding from 18 different *in vitro* studies (using the same “O1” operator sequence as the GROQ-seq assay)^20^. In an attempt to correct for variability across those studies, we normalized the LacI variant *K_d_* values by the wild-type *K_d_* within each study. Comparison with the log_g0 values from GROQ-seq has an approximately linear correlation with Spearman ⍴ = 0.65 and Pearson r = 0.62 (Figure 9D).

Recently, d’Oelsnitz et al. used a deep mutational scanning (DMS) assay similar to GROQ-seq to measure the sequence-function relationship for over 300,000 RamR variants^21^. They also reported a log_g0 function value (base-10 logarithm of uninduced transcription), enabling direct comparison with the GROQ-seq dataset reported here. For 3,305 RamR variants present in both datasets, that comparison shows a linear correlation (Spearman ⍴ = 0.68, Pearson r = 0.66) (Figure 9E).

We are not aware of any systematic mutagenesis data for VanR. However, Meyer et al., used directed evolution to engineer a variant of VanR with uninduced transcription approximately 100-fold lower than the wild-type^22,23^.Their improved variant had three mutations: T49I, A111V, and P179S. From our GROQ-seq dataset, the log_g0 values for single-mutant variants with those mutations are 1.49 ± 0.44, 3.05 ± 0.17, and 2.38 ± 0.20, respectively, compared with a wild-type log_g0 of 2.76 ± 0.08. Notably, two of these mutations (T49I and P179S) show reduced log_g0 relative to wild -type, broadly consistent with the improved repression phenotype reported by Meyer et al. In contrast, A111V exhibits near-wild-type activity, suggesting that its contribution to the triple-mutant phenotype may be context-dependent.

### T7 RNA Polymerase

T7 RNA polymerase (RNAP) is a widely used enzyme for producing massive quantities of mRNA and recombinant proteins at scale due to its exceptional transcriptional processivity and highly specific DNA promoter sequence recognition. Because of its broad utility, substantial research has focused on understanding T7 RNAP function and engineering more efficient variants. Despite this interest, deep mutational scanning of the enzyme has remained elusive, and mapping a highly detailed functional landscape on a per-residue resolution has not been achieved. The data presented here provide a foundation for engineering improved T7 RNAP variants and generating large, quality data to train future machine learning models for predicting function.

Although widely used, T7 RNAP is a difficult protein to study because of its toxicity to host cells^24^. For the T7 RNAP GROQ-seq, we implemented a simple plasmid design with constitutive expression of T7 RNAP and DHFR expression regulated by a T7 promoter (Supplemental 4A, see T7 RNAP Technical Bulletin for details^3^). In the data release^25^, the calibrated function value is the rate of transcription from the T7 promoter (log_g is the base-10 logarithm of that transcription rate). Since the circuit does not contain any inducible components, we ran the GROQ-seq assay with just two conditions: with and without TMP (Supplemental 4B). We then allocated the 24 available GROQ-seq conditions to run 12 identical replicates of the assay (12 samples with TMP and 12 samples without TMP). For the data release, we merged the barcode counting data across all 12 replicates and across multiple barcodes with the same amino acid sequence. To assess reproducibility (Figure 10B,C), we also analyzed the 12 replicates and individual barcodes separately. For T7 RNAP, we investigated single-site variant (SVL) and error-prone PCR multimutant libraries (epPCR).

#### T7 RNAP Variant Libraries

The GROQ-seq T7 RNAP data release includes sequence-function data for 34,641 unique variants. The SVL library contained 12,659 single-mutation variants, representing 75.6% coverage of the 16,746 possible single-amino-acid variants (Table 2).

**Table 2:** T7 RNAP variant library statistics.

| Protein | Library Type | Total Unique Variants | Total Variants with Single Mutation | Total Possible Designs | Percent of total designs tested* | Mean Number of Barcodes Per Variant | Median Number of Mutations per Variant |
| --- | --- | --- | --- | --- | --- | --- | --- |
| T7 | epPCR | 20,148 | 1 | - | - | 1 | 13 |
|  | SVL | 14,492 | **12,658** | **16,746** | 75.6 | 1 | 1 |

\*Percentage of designs tested is calculated as the number of variants observed from that library divided by the total possible library space (bolded values).

For T7 RNAP, 31 positions were missing from the gene library synthesis used for the SVL libraries. Those missing positions are L37, E48, P100, T101, A102, P111, K164, N165, V166, K172, R173, V186, R215, C216, H230, Q324, A338, A413, K450, I489, L550, L582, L611, G645, D770, S776, V783, D812, S838, C839, and L842.

To assess biological reproducibility for T7 RNA polymerase variants, we compared measurements from independent barcodes corresponding to the same amino acid sequence variants within one of the 12 replicates in the GROQ-seq assay. That comparison shows moderately strong correlation with small error between barcode pairs (Figure 10A, Spearman ρ = 0.77, RMSD = 0.25). We also assessed reproducibility across replicates by comparing measurements of the same individual barcoded variants between all pairs of replicates. The Spearman correlation and RMSD metrics indicate consistency across those replicates (Figure 10B, C, Spearman ρ = 0.74-0.78, RMSD = 0.25-0.3) showing that GROQ-seq measurements are robust and yield reproducible functional estimates across replicates.

#### T7 RNAP GROQ-seq Functional Data

Compared to the GROQ-seq assays for transcription-factors and proteases, the T7 RNAP assay has noticeably lower dynamic range, with little or no ability to distinguish variants with higher activity than wild-type (Figure 11A, 12A).

**Figure 11:**
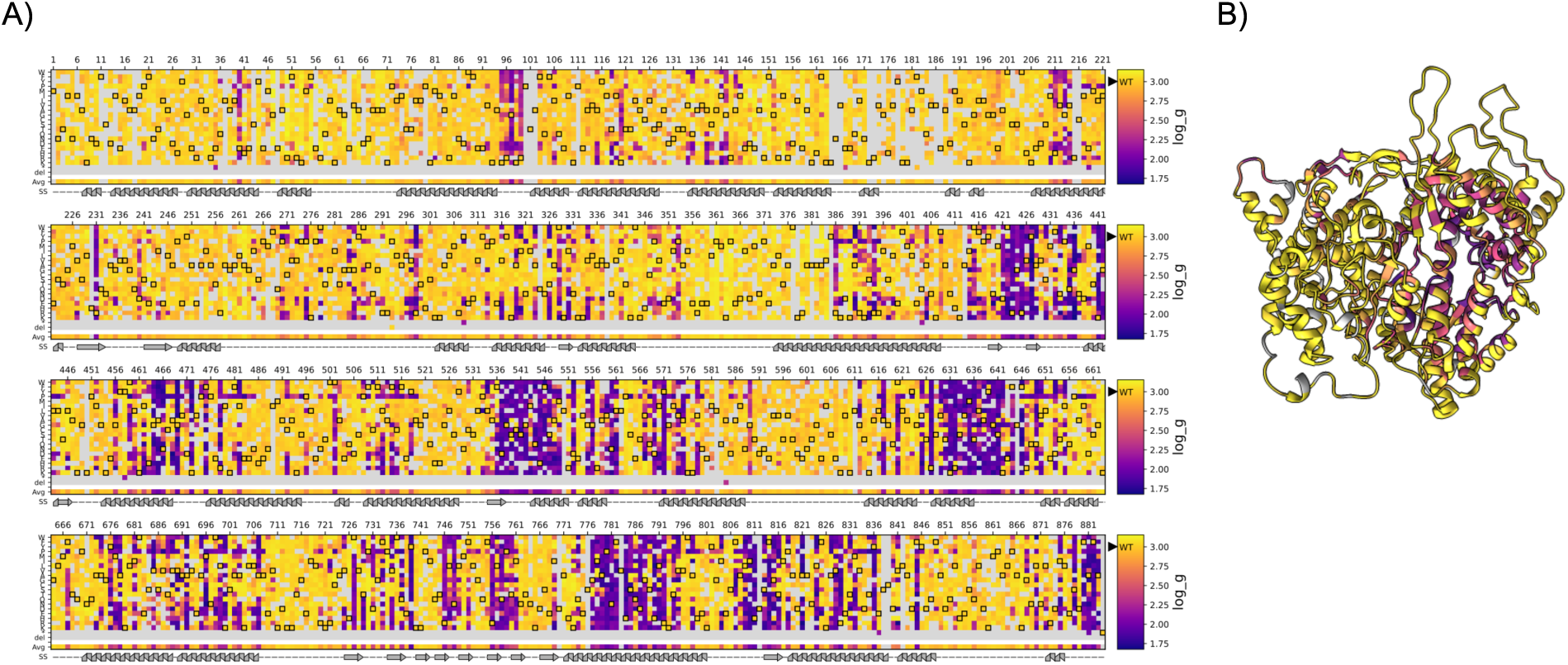
T7 RNAP substitution and deletion variant GROQ-seq data. (**A**) Heatmaps showing the function values for the base-10 logarithm of the rate of transcription from the T7 promoter single-mutant variants of T7 RNAP. The log_g values are expected to be directly related to the functional value of T7 RNAP, so lower log_g corresponds to lower T7 RNAP activity. The heatmap is split into four panels stacked vertically, with each subpanel spanning one-fourth the total length of the protein. The x-axis represents residue position, and the y-axis has the amino acids grouped by physicochemical properties of their side chains followed by the stop codon (*), deletion (del), and the average effect of mutations at each position (avg). Each cell is colored on a continuous scale spanning the observed range of function values shown on the right with the value of the wild-type marked by the filled black triangle labeled “WT”. The wild-type residue at each position is marked with a black outline in the heatmap. Missing data (e.g., missing variants from the library synthesis) and the average cell for positions with less than five reported substitutions are rendered in gray. A secondary structure annotation (helix, strand, loop) is drawn below each panel. (**B**) Protein structure with residues colored by the per-residue average log_g values for single amino acid substitutions (PDB: 1CEZ^26^) (with positions containing <5 mutations rendered in white). The DNA binding domain is oriented towards the top of the structure.

Like the GROQ-seq results for RamR, about half of all single substitutions in T7 RNAP result in activity similar to or greater than wild-type (55.4%; Figure 12B, where similar to or greater than wild-type is defined as within 2-fold of the wild-type activity or higher, i.e., greater than the wild-type log_g minus log_10_(2)). T7 RNAP is predominantly alpha-helical (50%), which would predict higher mutational robustness, based on the results of von der Dunk et al., and the data bear this out^16^.

**Figure 12:**
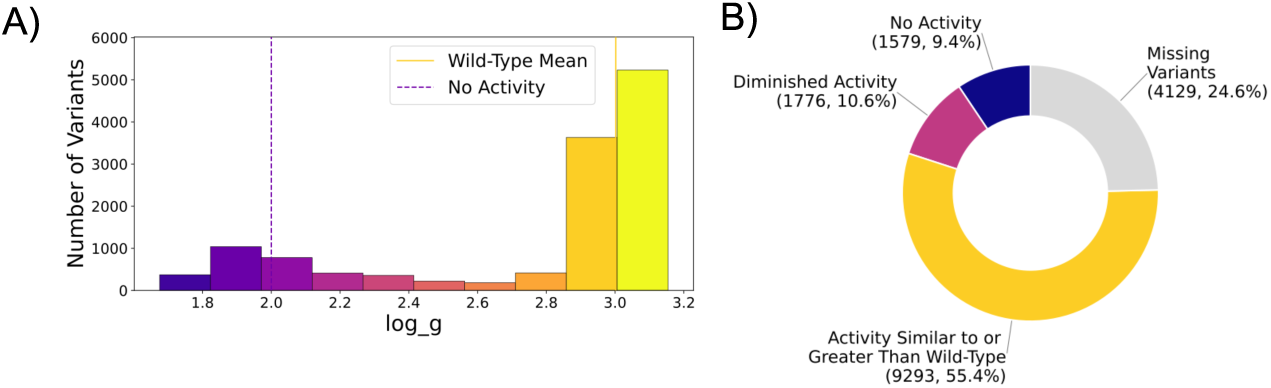
Comparison of T7 RNAP single residue substitution and deletion variants to wild-type function. (**A**) The histogram plot shows the range of the measured outcomes split into 10 equal-sized intervals on the x-axis and the number of variants sampling each interval on the y-axis. Each bar is colored using the same colormap as the corresponding heatmap in Figure 11. The wild-type mean is indicated by the solid line; the threshold for the no-activity category (dashed line) is set based on the lowest functional value for variants in the calibration ladder; (**B**) The donut plot shows the proportion of amino acid substitutions with log_g values categorized relative to the wild-type: No activity (low log_g), diminished activity (log_g values between no-activity threshold and similar-to-wild-type activity), similar to or greater activity than wild-type (greater than two-fold minus the wild-type log_g). Missing variants are shown in gray.

Although T7 RNAP is very tolerant to mutations along most of its length (Figure 11A: high log_g values indicated by yellow in the heatmap), there are notable regions of mutational intolerance (visible as purple bands in the heatmap). Those intolerant regions are generally associated with structural features that are important for T7 RNAP function. For example, the first sharp stretch of poor mutational tolerance is a stretch of positively charged residues (K95, R96, K98, R99) involved in binding to the minor groove of the DNA^27^. Position R231, which is completely intolerant to mutations, is the first position in the beta-hairpin structure (R231 to E242) involved in transcript initiation^28^. The set of positions intolerant to mutations between D421 and T440 in the core-palm domain (K412 to G449)^29^ include R425, the conserved residue required for NTP recognition and binding^30^. The set of positions intolerant to mutations between F536 and M549 sit at the junction of the core-palm and palm-insertion (K450 to S527)^31^ domains and include D537, one of two aspartate residues critical for coordinating the Mg²⁺ ion at the active site. Positions R627, K631, and R632 in the O-helix form H-bonds with the incoming NTP^30,32^ and are intolerant to mutations as well. H784 and H811 are completely intolerant to substitutions. H784 contacts the 2’-OH group of the incoming NTP^33^, and H811 contacts the +1 guanine nucleotide of the T7 promoter needed for strong transcription^34^.

To measure the effects of multi-mutation combinations on the function for T7 RNAP, we included an epPCR library in the GROQ-seq measurement. Most T7 RNAP epPCR variants contained 8 to 17 mutations (Figure 13A). Consistent with expectations, T7 RNAP activity decreases with increasing mutational burden, with most variants containing 5 or more mutations exhibiting substantially reduced function (Figure 13B). In contrast to the TF and TEV protease datasets, we obtain hundreds of measurements for T7 RNAP variants with >20 mutations, extending into a rarely sampled high-mutation regime.

**Figure 13:**
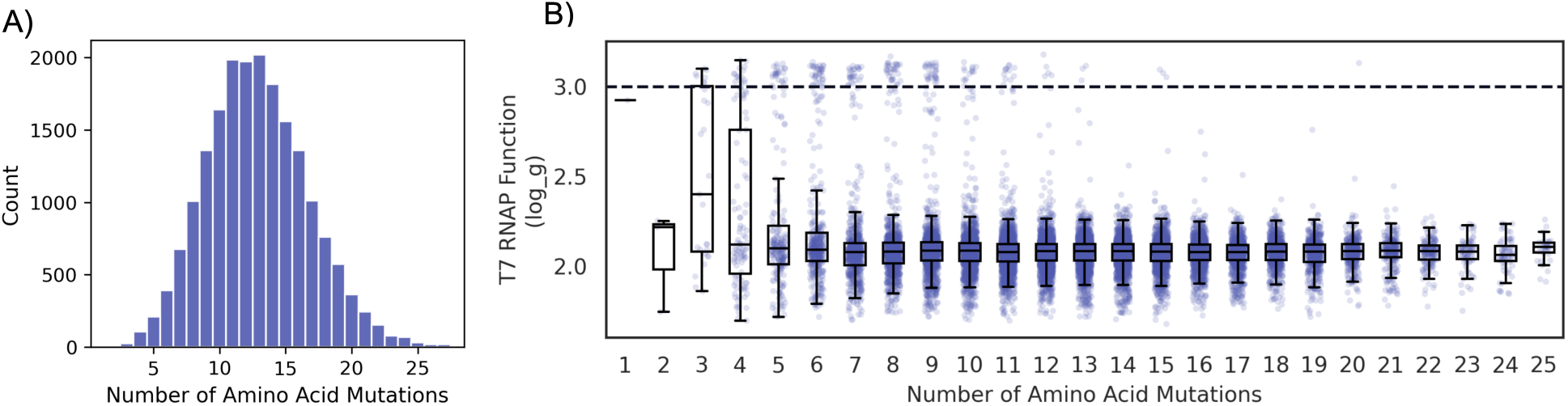
T7 RNAP epPCR library variant function. (**A**) Distribution of number of variants; (**B**) Functional values for epPCR variants, dashed-line represents wild-type level activity, whiskers indicate ±1.5× inter-quartile range.

#### Comparison to Existing Datasets

While we are unable to find previous studies with comprehensive T7 RNAP mutagenesis data, several site-specific mutagenesis studies have investigated T7 RNAP variants on a smaller scale to understand transcriptional activity beyond the role of the active sites. Gardner et al.^35^ quantified the kinetic parameters (*k_cat_* and *K_m_*) of several variants with substitutions to the C-terminal FAFA peptide loop of wild-type T7. Although data are limited, comparison between their results for catalytic efficiencies (*k_cat_*/*K_m_*) and log_g from the GROQ-seq assay shows a clear correlation (Figure 14).

**Figure 14:**
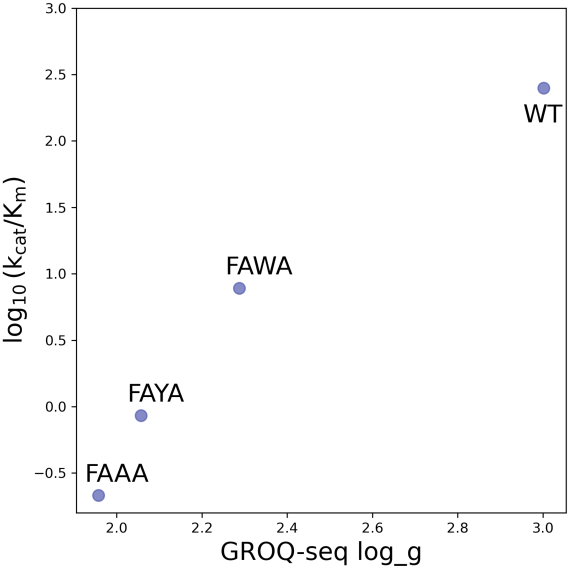
Comparison of GROQ-seq function data to k_cat_/K_m_ from Gardner et al. The text adjacent to each data point is the C-terminal 4-amino acid peptide for the T7 RNAP variant; “WT” indicates the wild-type, with C-terminal FAFA.

### TEV Protease

Tobacco Etch Virus (TEV) protease, in its natural context, provides an essential mechanism for viral polyprotein processing and replication. It uniquely recognizes the seven–amino acid sequence ENLYFQS and cleaves between Q and S. Because of its high specificity and reliability in protein processing applications, TEV protease has become one of the most widely used proteases in biotechnology. It has been extensively engineered in laboratory settings to improve catalytic efficiency, solubility, and target specificity^36–39^. However, no comprehensive DMS study, with mutations applied across the entire TEV protease protein, has been previously reported.

For the GROQ-seq protease platform, we designed a plasmid with two individually inducible expression cassettes: one controlling expression of the TEV protease variants, and a second controlling expression of a split DHFR reporter^40^ (Supplemental Figure 5A, see TEV Protease Technical Bulletin for details^4^). The two halves of the split DHFR are connected by a flexible peptide linker that contains the TEV protease substrate sequence (ENLYFQS). In the presence of TMP, the fitness of the host cells is modulated by the level of intact DHFR-substrate. Active TEV protease variants cleave the linker, reducing the level of intact DHFR-substrate, resulting in decreased fitness.

To explore the multi-dimensional space of TEV protease function and biophysics, we implemented the GROQ-seq assay as a multi-parametric measurement using a combination of ten different induction levels of TEV protease expression and two different induction levels of DHFR-substrate expression (Supplemental Figure 5B). This is similar to conventional *in vitro* enzyme kinetics assays that use different enzyme and/or substrate concentrations to extract the enzyme kinetic parameters (*k_cat_*, *K_m_*). For the current data release^41^, the calibrated function values are the level of intact DHFR (i.e., substrate that was not cleaved by protease activity) across 20 different samples (ten protease levels x two DHFR-substrate levels) (Supplemental Figure 5C). Here, to give an initial overview of the data, we focus on results for the lower induction level of DHFR and the highest induction level of TEV protease, the log_dhfr_S12 column from the released data tables, which is the base-10 logarithm of the level of intact DHFR substrate for that condition. Those results show the largest range of functional outcomes across the measured TEV protease variants (Supplemental Figure 5C). The level of intact DHFR substrate is inversely related to the catalytic activity of the protease, so lower log_dhfr_S12 values correspond to higher activity.

Below we present TEV protease, where we investigated an SSVL and epPCR library.

#### TEV Protease Variant Libraries

The GROQ-seq TEV protease data release includes sequence-function data for 18,465 variants. The SSVL library contains 8,623 single-mutation variants, representing 91.2% coverage of the 9,460 possible single-amino-acid variants. (Table 3)

**Table 3:** TEV Protease Variant Library Statistics.

| Protein | Library Type | Total Unique Variants | Total Variants with Single Mutation | Total Possible Designs | Percent of total designs tested* | Mean Number of Barcodes Per Variant | Median Number of Mutations per Variant | Mean Number of Mutations per Variant |
| --- | --- | --- | --- | --- | --- | --- | --- | --- |
| TEV Protease | epPCR | 9,407 | 454 | - | - | 1 | 4 | 5 |
|  | SSVL | 9,788 | **8,623** | **9,460** | 91.2 | 3 | 1 | 1.1 |

*\*Percentage of designs tested is calculated as the number of variants observed from that library divided by the total possible library space (bolded values).*

Some positions were missing from the gene library synthesis used for the SSVL libraries. Those missing positions are G36 and N52.

To assess biological reproducibility, two barcode measurements were randomly sampled for each amino acid variant following the analysis method described in Spinner et al.^1^. Measurements were highly consistent (RMSD = 0.15), with strong correlations (Spearman ρ = 0.90; Figure 15A). Protocol reproducibility was assessed by repeating the GROQ-seq assay on two separate days using the TEV protease SSVL library. Measurements per variant were highly consistent across runs (RMSD = 0.11) with very strong correlation (Spearman ρ = 0.95; Figure 15B). These results demonstrate robust reproducibility across barcodes and across distinct regions of the assay’s dynamic range along with highlighting GROQ-seq’s reproducibility across independent experimental replicates.

**Figure 15:**
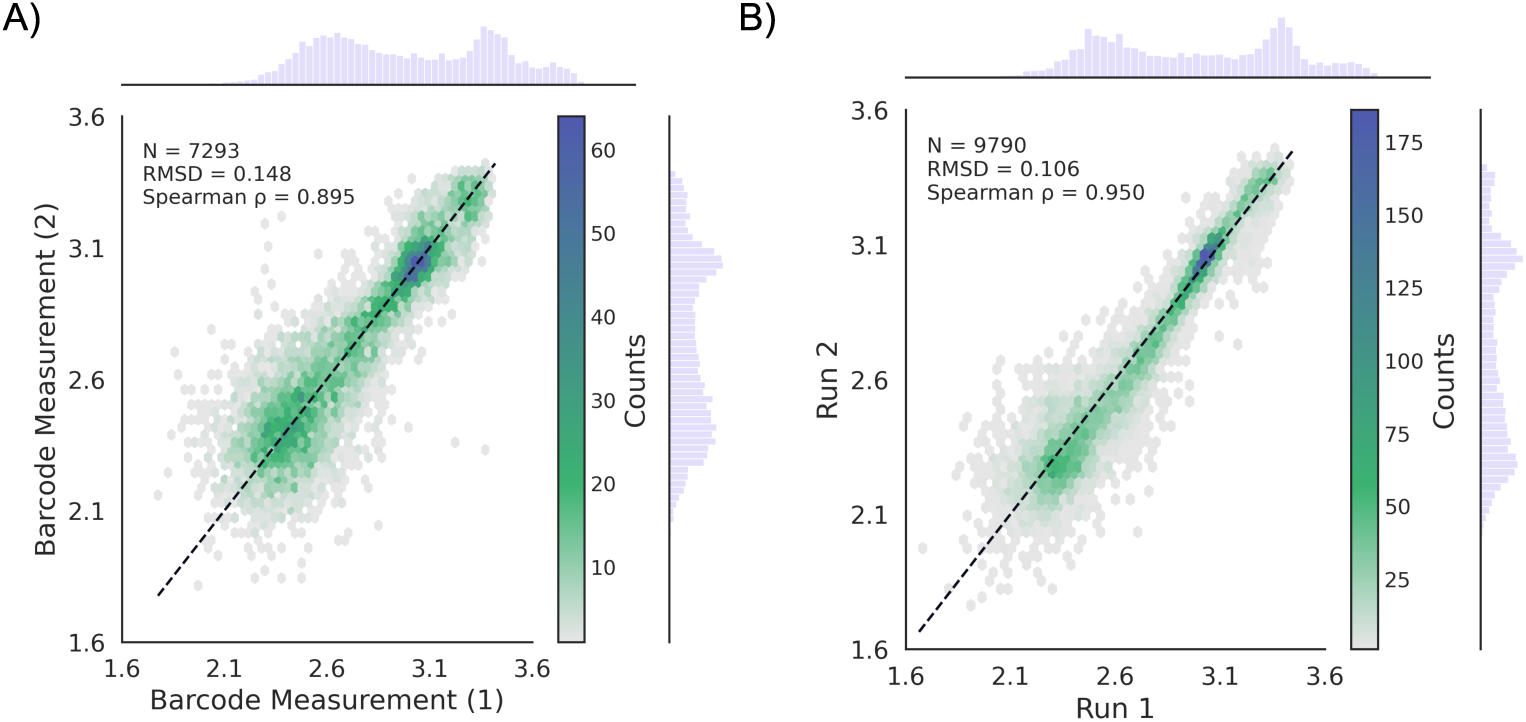
TEV protease biological and protocol reproducibility. (**A**) Comparing two barcodes per variant; (**B**) Comparing results from two separate runs per variant

#### TEV Protease GROQ-seq Functional Data

For single-mutant TEV protease variants, the GROQ-seq results are shown in Figure 16 and 17 for substitutions and deletions and Figure 18 for insertions. The components of each figure are similar to the TF and T7 RNAP figures: heatmaps summarize position-specific functional effects, protein structures visualize spatial patterns, histograms describe the distribution of single-mutant function values, and donut plots show the proportion of functional categories across the library.

**Figure 16:**
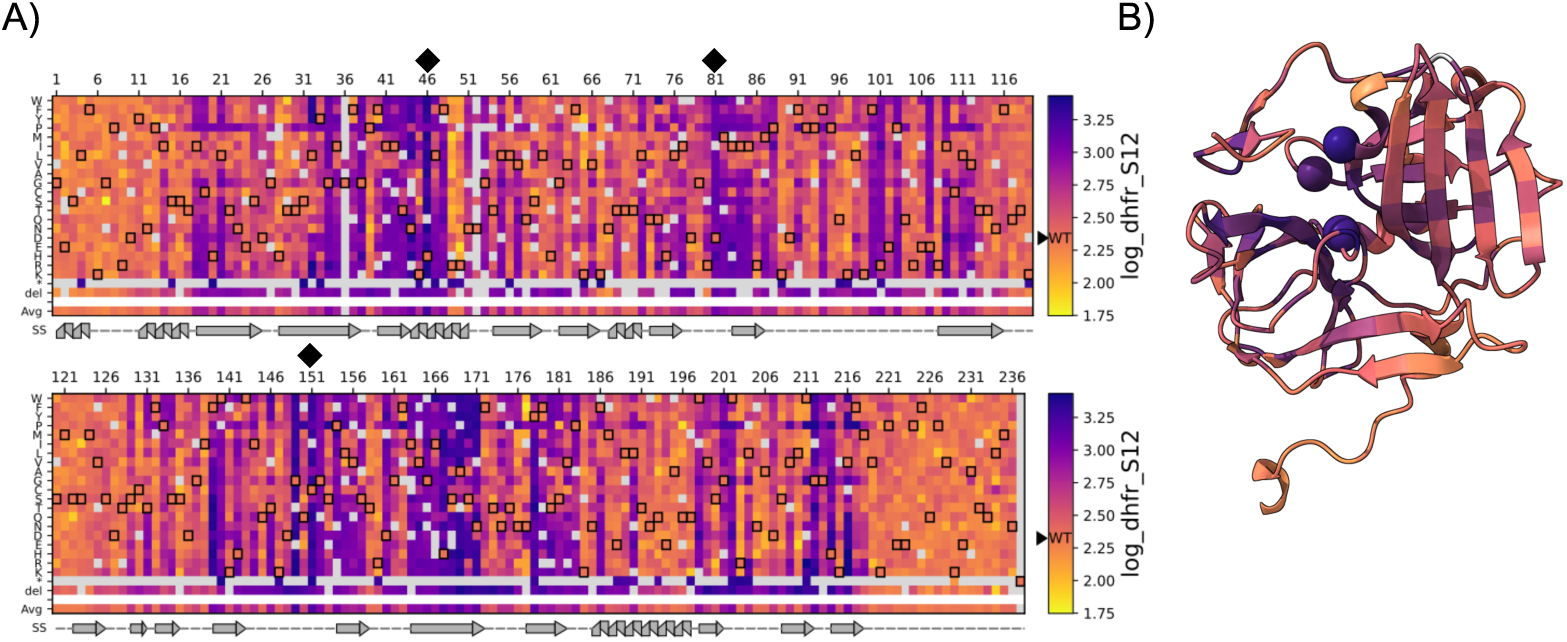
TEV protease single residue substitution and deletion variants GROQ-seq data. (**A**) Heatmap showing the function values for the log_dhfr_S12 column from the released data tables for the single mutant-variants of TEV protease: base-10 logarithm of the level of intact DHFR substrate for the GROQ-seq condition with the lower induction level of DHFR-substrate and the highest induction level of TEV protease. The level of intact DHFR substrate is inversely related to the catalytic activity of the protease, so lower log_dhfr_S12 values correspond to higher activity. The heatmap is split into two panels stacked vertically, with each subpanel spanning half the total length of the protein.The x-axis represents residue position, and the y-axis has the amino acids grouped by physicochemical properties of their side chains followed by the stop codon (*), deletion (del), and the average effect of mutations at each position (avg). Each cell is colored on a continuous scale spanning the observed range of function values shown on the right with the value of the wild-type marked by the filled black triangle labeled “WT”. The wild-type residue at each position is marked with a black outline in the heatmap. Missing data (e.g., missing variants from the library synthesis), and the average cell of positions with less than 5 reported substitutions are rendered in gray. A secondary structure annotation (helix, strand, loop) is drawn below each panel. Black diamonds denote catalytic residues (H46, D81, and C151); (**B**) Protein structure images with residues colored by the average log_dhfr_S12 values for single amino acid substitutions (TEV PDB: 1LVM^42^). One monomer of the biological assembly is colored by the per-residue average functional value (with positions containing <5 mutations rendered in white) with the residues forming the catalytic triad indicated by spheres (H46, D81, and C151).

**Figure 17:**
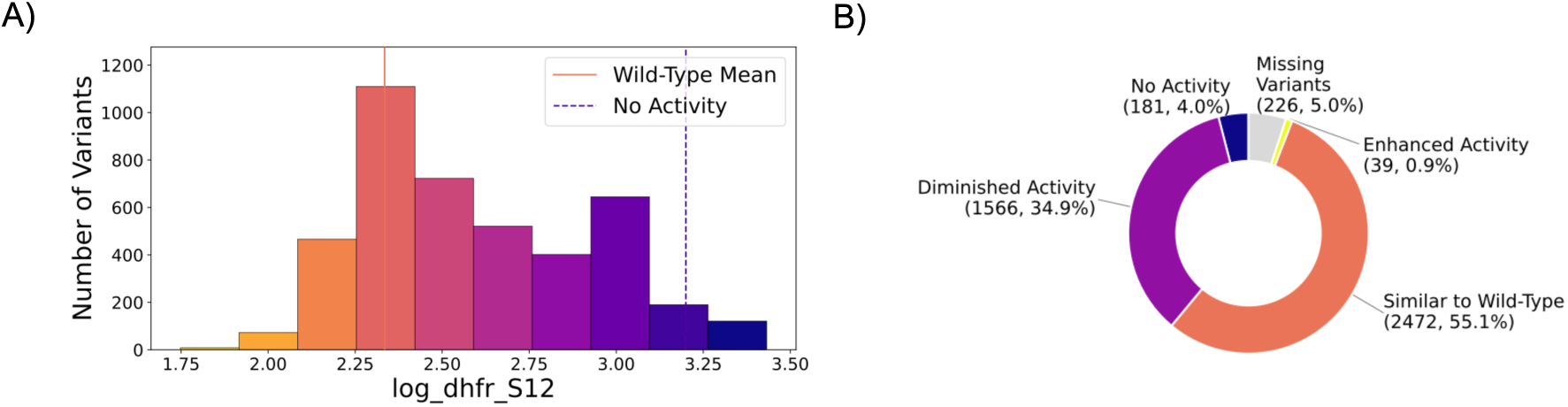
Comparison of TEV protease single residue substitution and deletion variants to wild-type function. (**A**) The histogram plot shows the range of the measured outcomes split into 10 equal-sized intervals on the x-axis and the number of variants sampling each interval on the y-axis. Each bar is colored using the same colormap as the corresponding heatmap in Figure 16. The wild-type mean is indicated by the solid line; the threshold for the no-activity category is set to the average of the log_dhfr_S12 values of the stop codons for the first 100 residues since this will not include the catalytic triad residues. The colors of both the lines correspond to their respective values in Figure 16A; (**B**) The donut plots shows the proportion of single substitutions with log_dhfr_S12 values categorized relative to the wild-type: No activity (high log_dhfr_S12), diminished activity (log_dhfr_S12 values between no-activity threshold and similar-to-wild-type), similar to wild-type (within 2-fold of wild-type mean on either direction), stronger than wild-type (low log_dhfr_S12). Missing variants are shown in gray.

**Figure 18:**
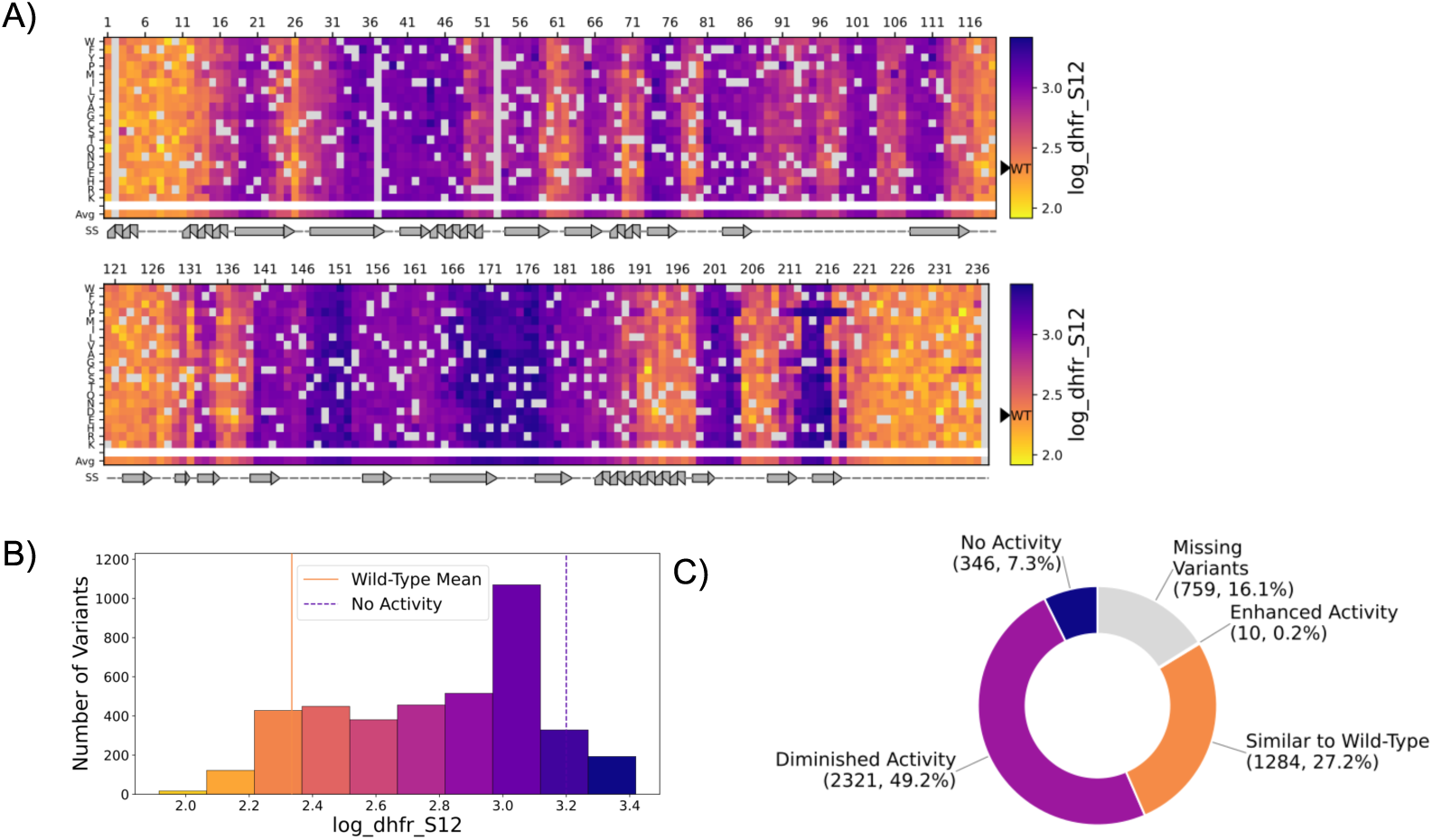
TEV Protease single residue insertion variants GROQ-seq data. (**A**) Heatmap showing the function values for the log_dhfr_S12 column from the released data tables for the single insertion-variants of TEV protease: base-10 logarithm of the level of intact DHFR substrate for the GROQ-seq condition with the lower induction level of DHFR-substrate and the highest induction level of TEV protease. The level of intact DHFR substrate is inversely related to the catalytic activity of the protease, so lower log_dhfr_S12 values correspond to higher activity. The heatmap is split into two panels stacked vertically, with each subpanel spanning half the total length of the protein.The x-axis represents residue position, and the y-axis has the amino acids grouped by physicochemical properties of their side chains followed the average effect of insertions at each position (avg). Each cell is colored on a continuous scale spanning the observed range of function values shown on the right with the value of the wild-type marked by the filled black triangle labeled “WT”. Missing data (e.g., missing variants from the library synthesis), and the average cell of positions with less than 5 reported substitutions are rendered in gray. A secondary structure annotation (helix, strand, loop) is drawn below each panel. (**B**) The histogram plot shows the range of the measured outcomes split into 10 equal-sized intervals on the x-axis and the number of variants sampling each interval on the y-axis. Each bar is colored using the same colormap as the corresponding heatmap in Figure 18A. The wild-type mean is indicated by the solid line; the threshold for the no-activity category is set to the average of the log_dhfr_S12 values of the stop codons for the first 100 residues since this will not include the catalytic triad residues. The colors of both the lines correspond to their respective values in Figure 18A; (**B**) The donut plots shows the proportion of single insertions with log_dhfr_S12 values categorized relative to the wild-type: No activity (high log_dhfr_S12), diminished activity (log_dhfr_S12 values between no-activity threshold and similar-to-wild-type), similar to wild-type (within 2-fold of wild-type mean on either direction), stronger than wild-type (low log_dhfr_S12). Missing variants are shown in gray.

Consistent with the findings for transcription factors, a large fraction of substitutions (56%) result in protease activity similar to or greater than wild-type (where similar to wild-type is defined as within 2-fold of the wild-type TEV protease activity, i.e., within ± log_10_(2) of the wild-type value for log_dhfr_S12) (Figure 17A, 18B). In contrast to the transcription factors, the structure of TEV has low alpha-helical content (14%) and is dominated by beta-sheet (39%). Thus, based on the findings of von der Dunk et al., we would expect TEV to have lower tolerance to mutations. However, our results contradict this expectation. This observation could also be attributed to the differences in the enzyme vs. DNA-binding protein functional requirements.

Although the fraction of substitutions that enhance function is much lower for TEV protease than the transcription factors, there are still notable positions where multiple different substitutions increase TEV protease activity.

At position N177, large hydrophobic substitutions (F, Y, W, M) give the highest activities (near 2-fold to over 3-fold decrease in log_dhfr_S12 relative to wild-type). Large positively charged substitutions (R, K, H) at position N177 give a smaller increase in activity (between 1.4-fold and 1.7-fold; Figure 16A).

Position R49 shows a broad set of activity-enhancing substitutions, with the highest activities resulting from M, A, T, and V substitutions (greater than 2-fold decrease in log_dhfr_S12 relative to wild-type), but significant increases in activity also for I, C, W, Q, L, Y, N, and S substitutions (Figure 16A). Several substitutions at position R50, immediately adjacent to R49, also increase protease activity, but with a distinct pattern of side-chain preference: R50E has a nearly 3-fold decrease in log_dhfr_S12 relative to wild-type, and substitutions to H, W, Y, T, and F also result in significant activity enhancement. (Figure 16A)

Unsurprisingly, the catalytic triad H46, D81, and C151 are particularly intolerant to a broad range of substitutions, consistent with their essential role in catalysis (Figure 16A). Furthermore, positions F139 and W140 appear to tolerate some hydrophobic residues but are intolerant to most other substitutions. This is consistent with the role of those positions in the S4 pocket of TEV protease in making hydrophobic interactions with the leucine in the substrate peptide^42^.

The other substrate-interacting residues that are intolerant to substitutions include A169, which also forms hydrophobic interactions with the leucine in the substrate. Notably, T170 and N171, which are adjacent to A169, also tolerate substitutions poorly^42^ (Figure 16A).

G212 is extremely sensitive to substitutions and only tolerates glycine. This is explained by its role in H-bond formation with the substrate’s threonine^42^.

Positions F100 are R101 intolerant to substitutions as well. However, there are no known interactions reported for these positions with the substrate, and they are approximately 23 Å away from the substrate. In an MSA reported by Phan et al., these positions appear highly non-conserved across other proteins with similar fold, and they fall on a loop on the structure^42^ .

In TEV protease, as with the TFs, regions that tolerate insertions are similar to regions that tolerate substitutions (Figure 18A). However, unlike the TFs, in TEV protease, the positions where substitutions increase activity do not have corresponding regions where insertions also increase activity. For example, all insertions near R49 and R50 decrease activity, and insertions near N177 generally result in very low or no detectable activity. More generally, residues that are part of beta-sheets in one of the two beta-barrel structures generally do not tolerate insertions.

To examine the effects of multi-mutation combinations on the function for TEV protease, we included an epRCR library in the GROQ-seq measurement. Most variants in the TEV protease epPCR library contained between 2–5 mutations, with a tail extending to higher mutation counts (Figure 19A). TEV protease activity generally decreases (i.e. intact DHFR increases, as seen in the plot) as the number of mutations increase, reflecting the cumulative disruptive effects of multiple amino acid substitutions (Figure 19B). Notably, there are a number variants with 4 or more mutations that still retain near-wild-type activity, indicating that TEV protease retains some tolerance to sequence perturbation despite being an enzyme with well-defined catalytic requirements. This is in agreement with the substitution and deletion variant data, where 56% of these variants retain nearly wild-type activity (Figure 17B).

**Figure 19:**
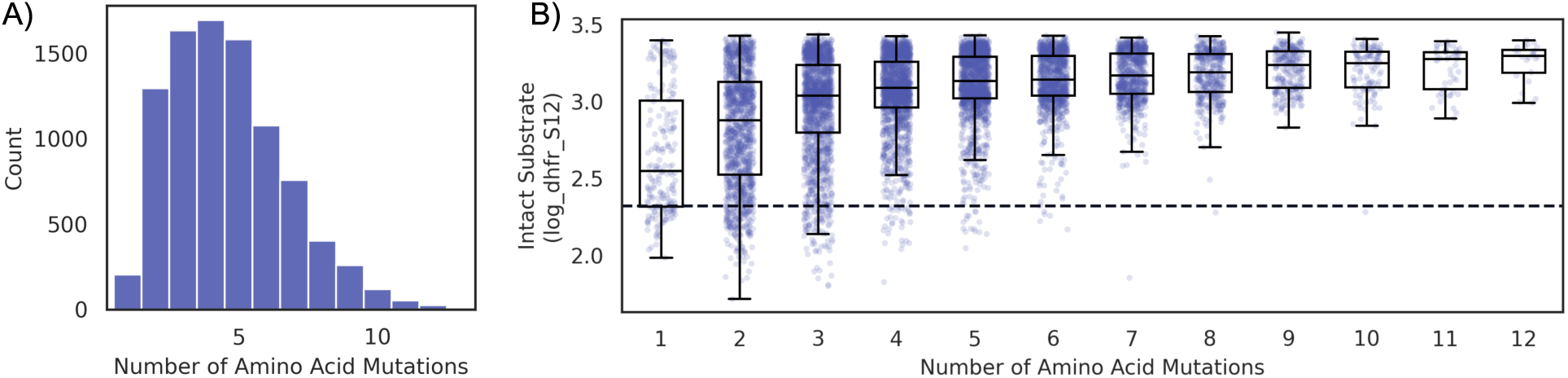
TEV protease epPCR GROQ-seq data. (**A**) Distribution of number of variants; (**B**) Functional values for epPCR variants, dashed-line represents wild-type level activity, whiskers indicate ±1.5× inter-quartile range.

#### Comparison to Existing Datasets

Sundar et al., recently reported a DMS dataset that includes results for 159,060 TEV protease variants (comprehensive four-position combinatorial mutagenesis)^43^. Of those, 75 single-mutant variants were shared with the GROQ-seq dataset reported here. Comparison between the Sundar et al. fitness scores and GROQ-seq log_dhfr_S12 results shows a non-linear correlation with Spearman ⍴ = -0.70 (Figure 20; negative correlation because lower values indicate higher TEV activity in GROQ-seq, whereas they indicate lower TEV activity in the Sundar et al. assay). The relationship between the Sundar et al. fitness scores and the GROQ-seq log_dhfr_S12 values seems to follow a sharp sigmoidal curve with most of the changes in fitness score occurring at high values of log_dhfr_S12. This is similar to the comparison between GROQ-seq and the DMS results reported by Meger et al. for LacI (Figure 9C), again indicating a wider dynamic range for the GROQ-seq measurement.

**Figure 20:**
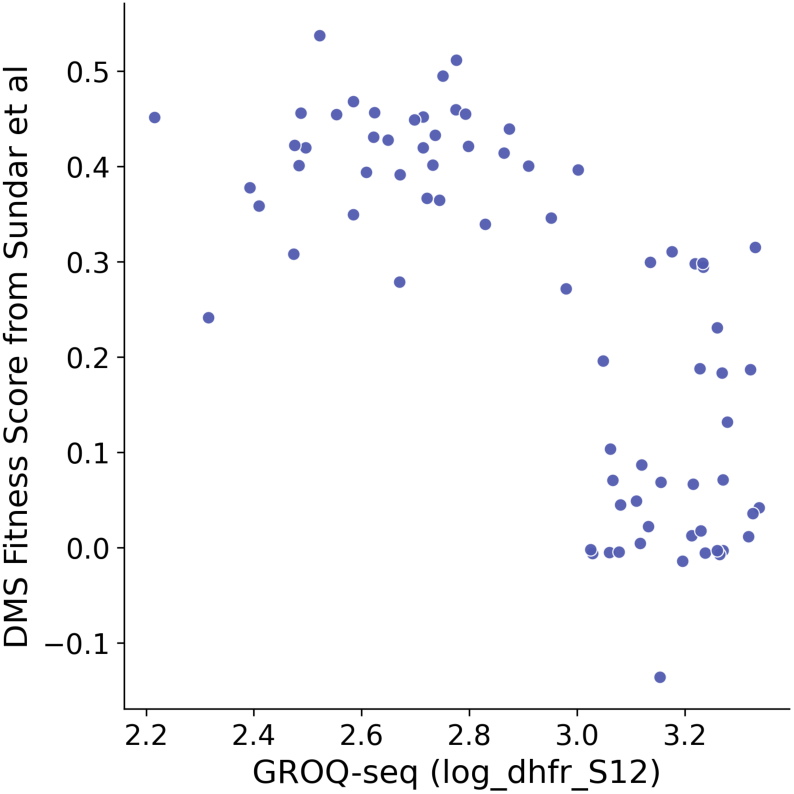
TEV Protease GROQ-seq data comparison to Sundar et al. dataset. Comparison of fitness score from Sundar et al and GROQ-seq functional data (Spearman = -0.70).

## Conclusion

The datasets presented here are the largest openly available DMS resources for these proteins. Among all three datasets presented, high reproducibility was observed between barcodes of the same variant, indicating robust experimental design and biological replicate reproducibility (Spinner et al.^1^,Figure 10, Figure 15).

The results show a surprising robustness to mutational perturbations: Across all five proteins for three different function types, between about 1/4 to 1/2 of all possible single amino acid substitutions result in protein activity similar to or greater than the wild-type protein (Figures 4A-C, Figure 12B, Figure 17B). Even for a large, multi-domain protein like T7 RNAP, with a complex, multi-step, multi-interaction function, over half of all possible substitutions retain wild-type-like activity. This contrasts with the conventional view that mutations generally degrade activity.

For mutations where the GROQ-seq results indicate decreased activity, the most deleterious often occur at positions that are known to be catalytic or otherwise essential for function, reinforcing the validity of our approach (e.g., TF Figures 1-3,5-7; T7 RNAP Figure 11; TEV protease Figures 16,18). These findings are further supported by agreement with previously published datasets (Figures 9, 14, 20).

In addition to delivering high quality data, collected and analyzed in a consistent manner for different protein types, each dataset also has unique value. The TF dataset is notable for its ability to compare three transcription factor families within a single calibrated framework, allowing detection of both increases and decreases in function including mutations that enhance TF-operator affinity. The TEV dataset stands out for its exceptional measurement resolution and dynamic range, enabled by testing across 20 assay conditions that capture multidimensional biophysical effects. Meanwhile, the T7 RNAP dataset represents a major technical achievement due to the sheer size of the protein. To our knowledge, this is the only deep mutational scanning effort on this protein to date. Together, these datasets provide a comprehensive and high-quality view of sequence-function relationships across these three diverse protein functions, pointing toward a future where protein function can be measured at scale with the rigor required for predictive biology.

## Disclaimer

Certain commercial equipment, instruments, or materials are identified in this paper in order to specify the experimental procedure adequately. Such identification is not intended to imply recommendation or endorsement by the National Institute of Standards and Technology, nor is it intended to imply that the materials or equipment identified are necessarily the best available for the purpose.

## Funding Statement

The Align Foundation is supported by Griffin Catalyst and Schmidt Sciences.

The DAMP Lab acknowledges laboratory support from its industrial members including Hamilton, OpenTrons, MITRE, SciSure, New England BioLabs, Transfyr, and SiLA.

S.D acknowledges funding from the Harvard Medical School Synthetic Biology HIVE. E.D. acknowledges support by the Francis Crick Institute which receives its core funding to E.D. from Cancer Research UK (CC2239), the UK Medical Research Council (CC2239), and the Wellcome Trust (CC2239).

## Supplemental Information

### GROQ-seq Experiment

Barcoded protein variant libraries are grown in pooled culture across timepoints under defined induction and selection conditions, with plasmid DNA extracted after each growth phase to capture changes in variant abundance (Supplemental Figure 1). These measurements are converted into quantitative functional readouts through barcode sequencing and internal calibration, enabling high-throughput, mapping of sequence–function relationships^1^.

**Supplementary Figure 1:**
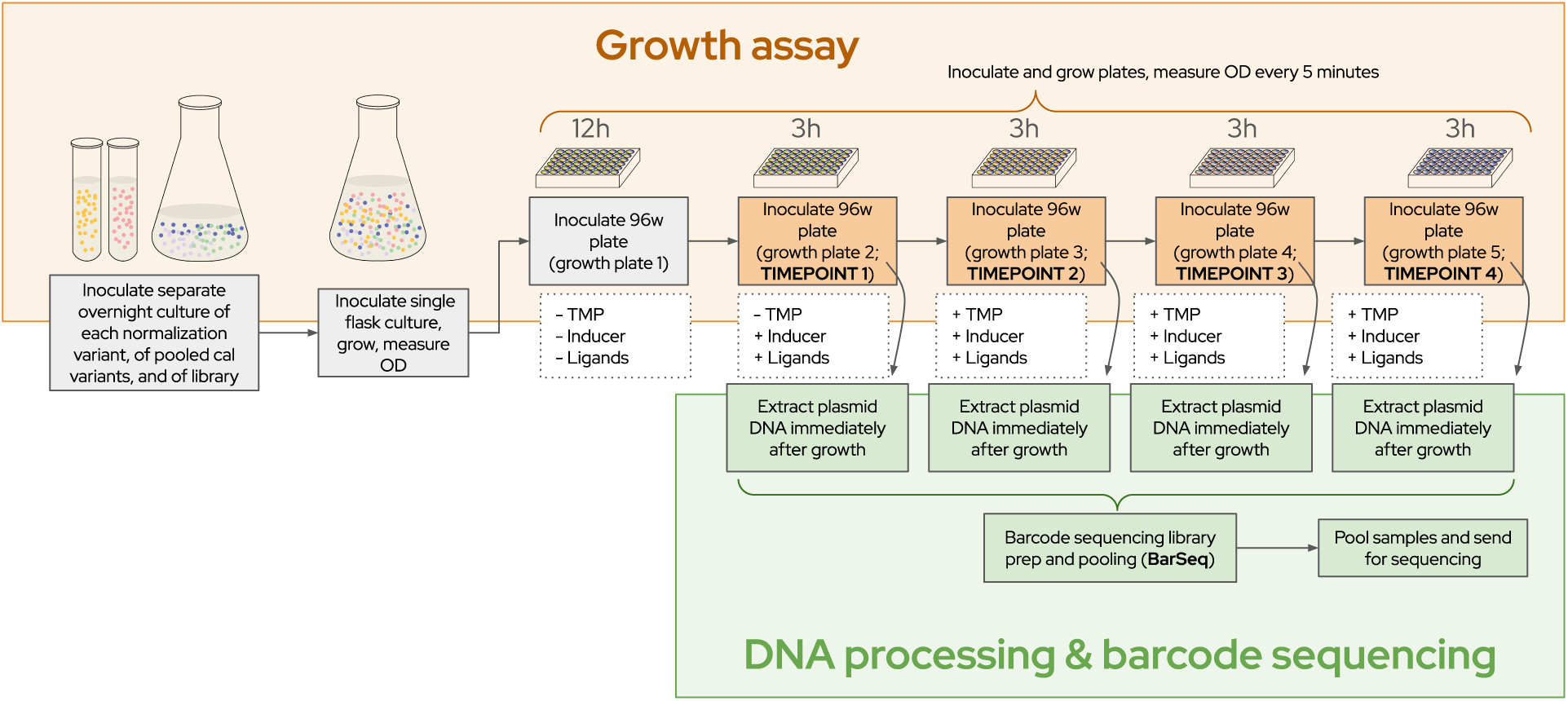
Schematic overview of the GROQ-seq growth assay and sequencing workflow.

### GROQ-seq Analysis Pipeline

GROQ-seq plasmid libraries contain protein variants coupled to unique barcodes. After each growth timepoint, cells are harvested, the barcode region is amplified from extracted plasmids and sequenced via Illumina short-read sequencing. The Bartender analysis workflow (Supplemental Figure 2: Bartender) clusters these reads into unique barcode identities (using bartender1.1^44^, plus extensions^45^, removing noise and producing a count table that records how many times each barcode has been detected in each sample and at each time point. To map barcodes to their corresponding full plasmid sequences, Oxford Nanopore long-read sequencing is performed on the input library (Supplemental Figure 2: Nanopore long-read track). The long-read data links each barcode to its variant sequence, while the short-read data measures fitness across different conditions (e.g., with different TMP concentrations, with and without inducers or ligands).

Unlike typical deep mutational scanning approaches, GROQ-seq uses a protein function calibration ladder to convert raw fitness measurements into biologically interpretable function values (Supplemental Figure 2: Fitness → function calibration). For this calibration, we use Bayesian regression to map the barcode-sequencing-based fitness measurements across multiple experimental conditions onto a multi-parametric readout of protein function. The result is a dataset linking each unique protein sequence with one or more calibrated functional measurements with well-calibrated posterior uncertainties (Supplemental Figure 2 GROQ-Seq Analysis Pipeline).

After quality filtering and data cleaning, barcodes corresponding to the same protein sequence are identified and their short-read counts are pooled. The pooled counts are reanalyzed using a second-round of Bayesian regression, producing posterior estimates of function and uncertainty per protein variant (rather than per barcode). The final data table is a protein sequence to function dataset in which each amino acid sequence carries calibrated functional measurements with well-characterized uncertainties, directly linking genotype to phenotype (Supplemental Figure 2: Protein sequence to function dataset).

**Supplemental Figure 2:**
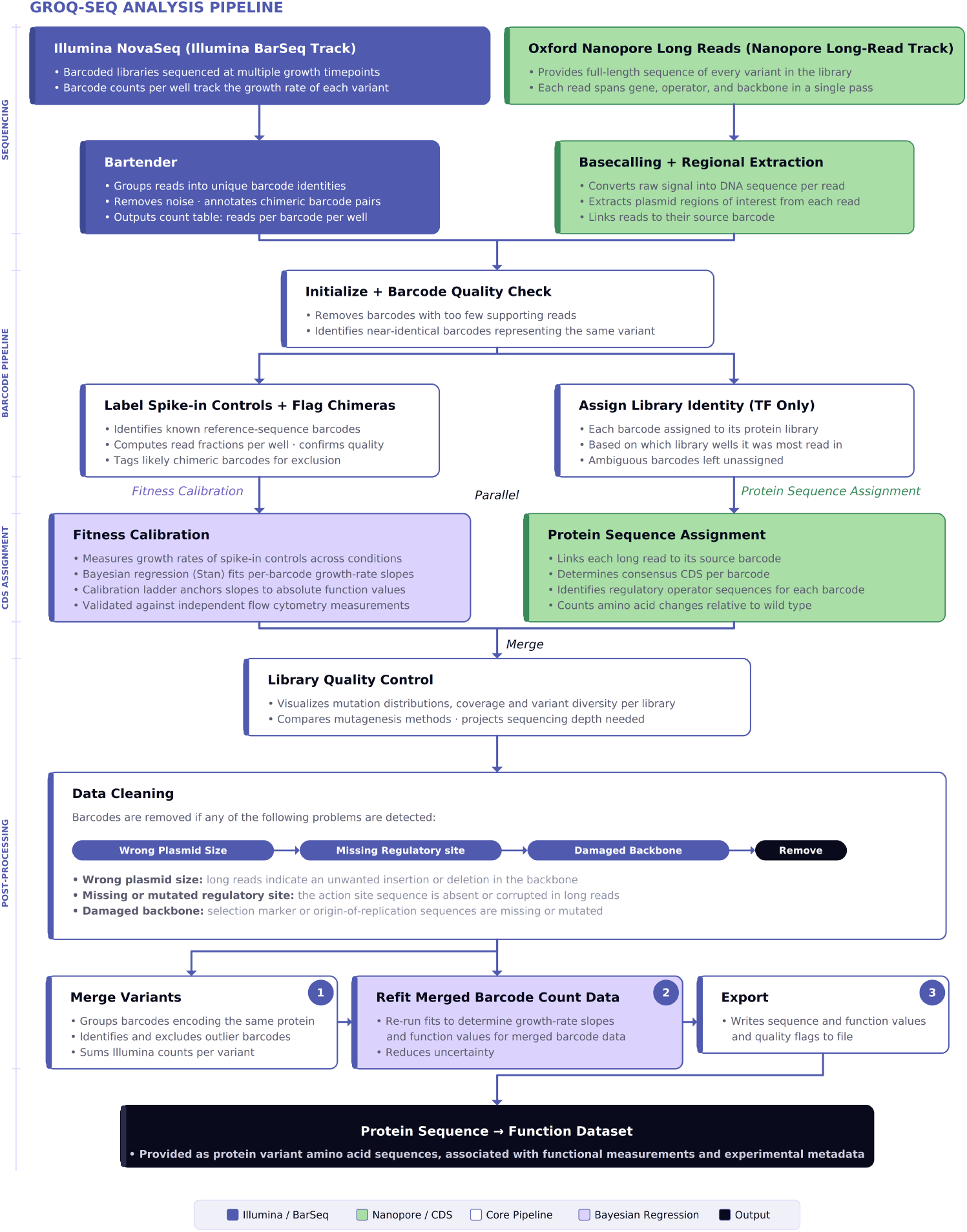
GROQ-seq Analysis Pipeline. Short-read barcode sequencing (Illumina NovaSeq, Illumina BarSeq Track) and full-plasmid long-read sequencing (Oxford Nanopore, Nanopore Long-Read Track) are conducted in parallel. Illumina reads are clustered into unique barcode identities by NISTBartender, producing a count table of reads per barcode per GROQ-seq sample. Barcodes with too few reads are discarded and near-identical barcodes are identified. Data for the calibration ladder plasmids is used to construct a calibration curve and Bayesian regression is used to convert from fitness to function values. In parallel, Nanopore reads are grouped by barcode, and the consensus from multiple Nanopore reads are used to determine the gene of interest (GOI) coding sequence and other sequence elements on the GROQ-seq plasmid. Barcodes failing quality checks are removed before barcodes with the same GOI amino acid sequence (i.e., same variant) are merged. Count data are refit, producing posterior estimates of function and uncertainty per protein variant, exported as a protein sequence → function dataset.

**Supplemental Figure 3:**
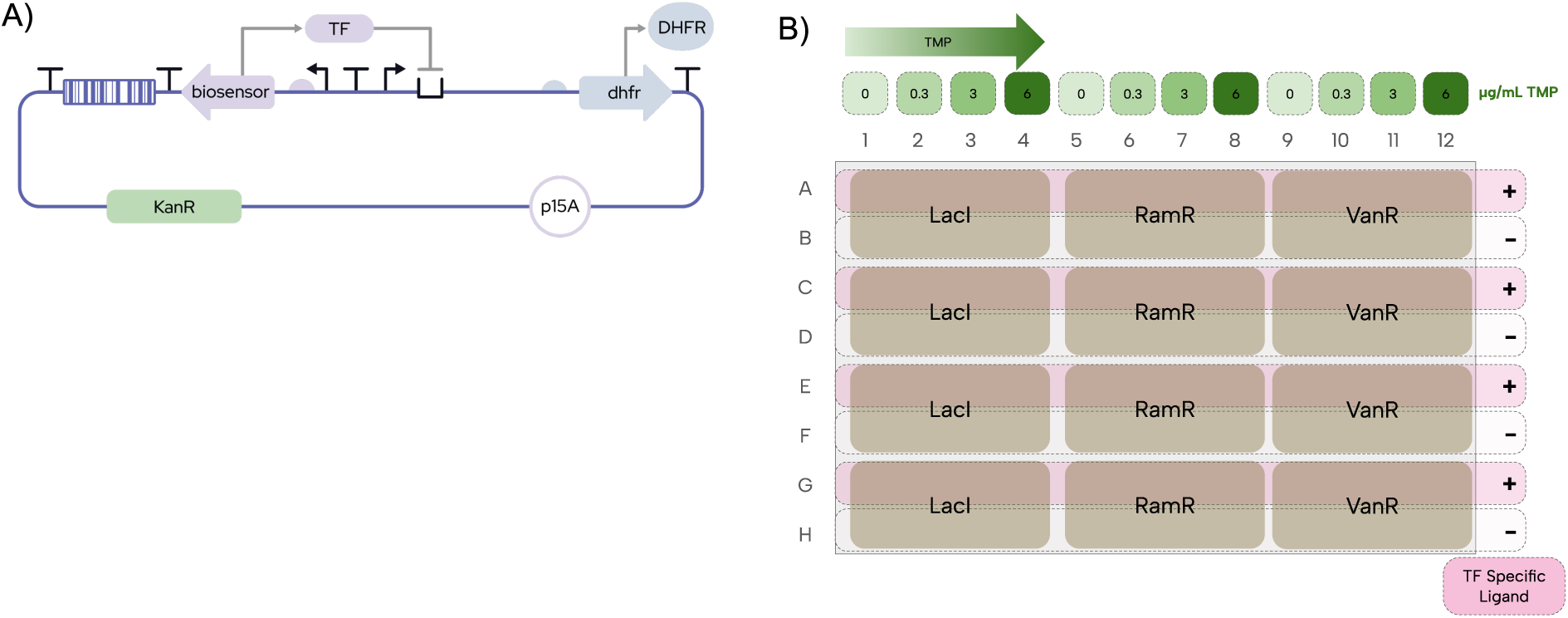
TF Experimental setup. (**A**) Schematic of plasmid circuit (TF=transcription factor, DHFR= dihydrofolate reductase,); (**B**) Schematic of plate layout used in GROQ-seq experiment (TMP=trimethoprim)

The plasmid circuit was designed so that the TF of interest regulates the expression of DHFR, a gene that provides resistance to the antibiotic trimethoprim (TMP). As a result, stronger repression (via binding of the transcription factor to its cognate DNA operator) leads to decreased resistance to TMP. Each variant plasmid tested in the experiment contained a unique random barcode, allowing for analysis across conditions (Supplemental Figure 3A).

Supplemental Figure 3B provides a summary of the plate layout used for the transcription factor GROQ-seq experiment. All variant libraries were tested across four TMP concentrations (0, 0.3 µg/mL, 3.0 µg/mL, and 6.0 µg/mL), in the presence and absence of cognate ligand (LacI with IPTG at 2000 µmol/L, RamR with 1S-TIQ at 250 µmol/L, and VanR with vanillic acid at 100 µmol/L). All variant libraries in each condition were tested in four replicate wells.

For each transcription factor investigated, three types of variant libraries were run: site saturation variant libraries (SSVL), site saturation mutagenesis (SSM), and error-prone PCR (epPCR). The SSM libraries specifically targeted known or hypothesized DNA-binding residues (positions Y17, Q18, and R22 for LacI; E42, G43, and R47 for RamR; and E33, R44, and M45 for VanR).

The dynamic range of the transcription factor assay was approximately 2.5 orders of magnitude.

The transcription factors LacI (UniProt ID: P03023; PDB: 1EFA), RamR (UniProt ID: A0A0F6AY66; PDB: 7N53), and VanR^23^ (AlphaFold model: AF-Q9A5Q5-F1; accession WP_010920250) were used in this study. Structural information was obtained from available crystallographic data or predicted models where experimental structures were not available.

**For more details on the experiment, please refer to the Technical Bulletin for GROQ-seq Transcription Factor Function Assay for details**^2^

**Supplemental Figure 4:**
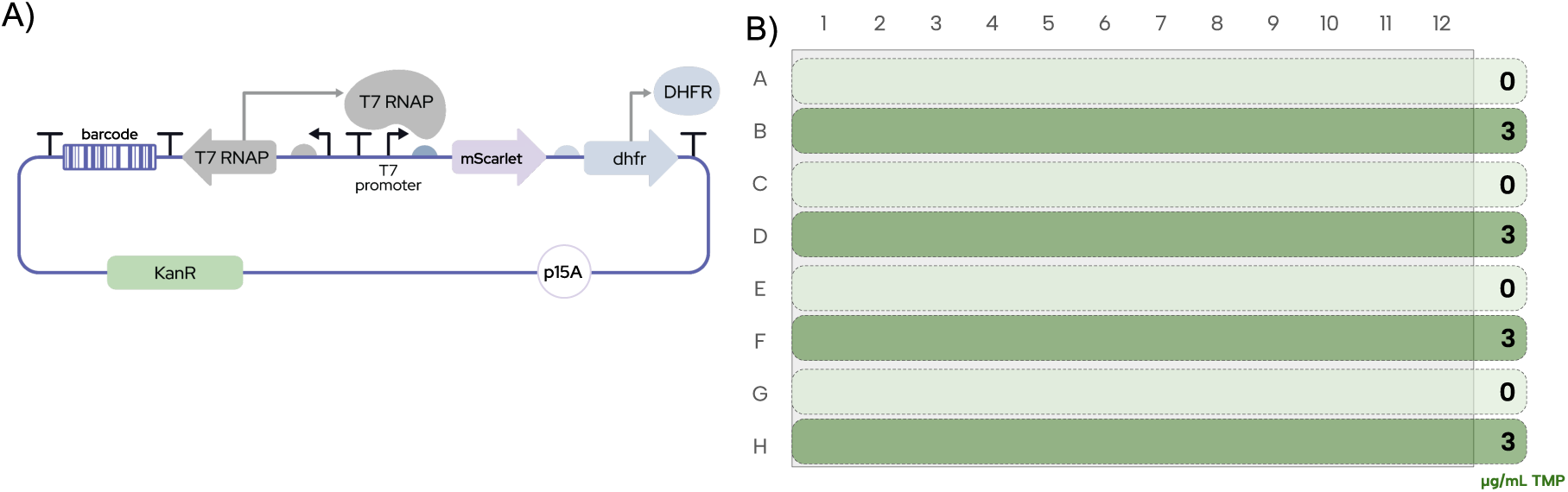
T7 RNAP Experiment Setup. (**A**) Barcoded T7 RNA polymerase (T7 RNAP) variants were constitutively expressed from plasmids containing a bicistronic selection cassette. This cassette includes a class III T7 promoter driving expression of both mScarlet and murine dihydrofolate reductase (mDHFR), coupling transcriptional activity to cellular fitness. Variants with higher functional activity confer increased fitness under trimethoprim (TMP) selection. (**B**) Plate conditions used in T7 RNAP GROQ-seq run

We used the GROQ-seq assay to measure the function of each variant in the single-site variant (SVL) and epPCR libraries for T7 RNAP with two two conditions (0 μg/mL and 3 μg/mL TMP).

The dynamic range of the T7 RNAP assay was approximately 0.7 orders of magnitude (∼5-fold).

The T7 RNAP structure used for Figure 18B was from PDB entry 1CEZ.

**For more details on the experiment, please refer to the Technical Bulletin for GROQ-seq T7 RNAP Function Assay for details**^3^

**Supplemental Figure 5:**
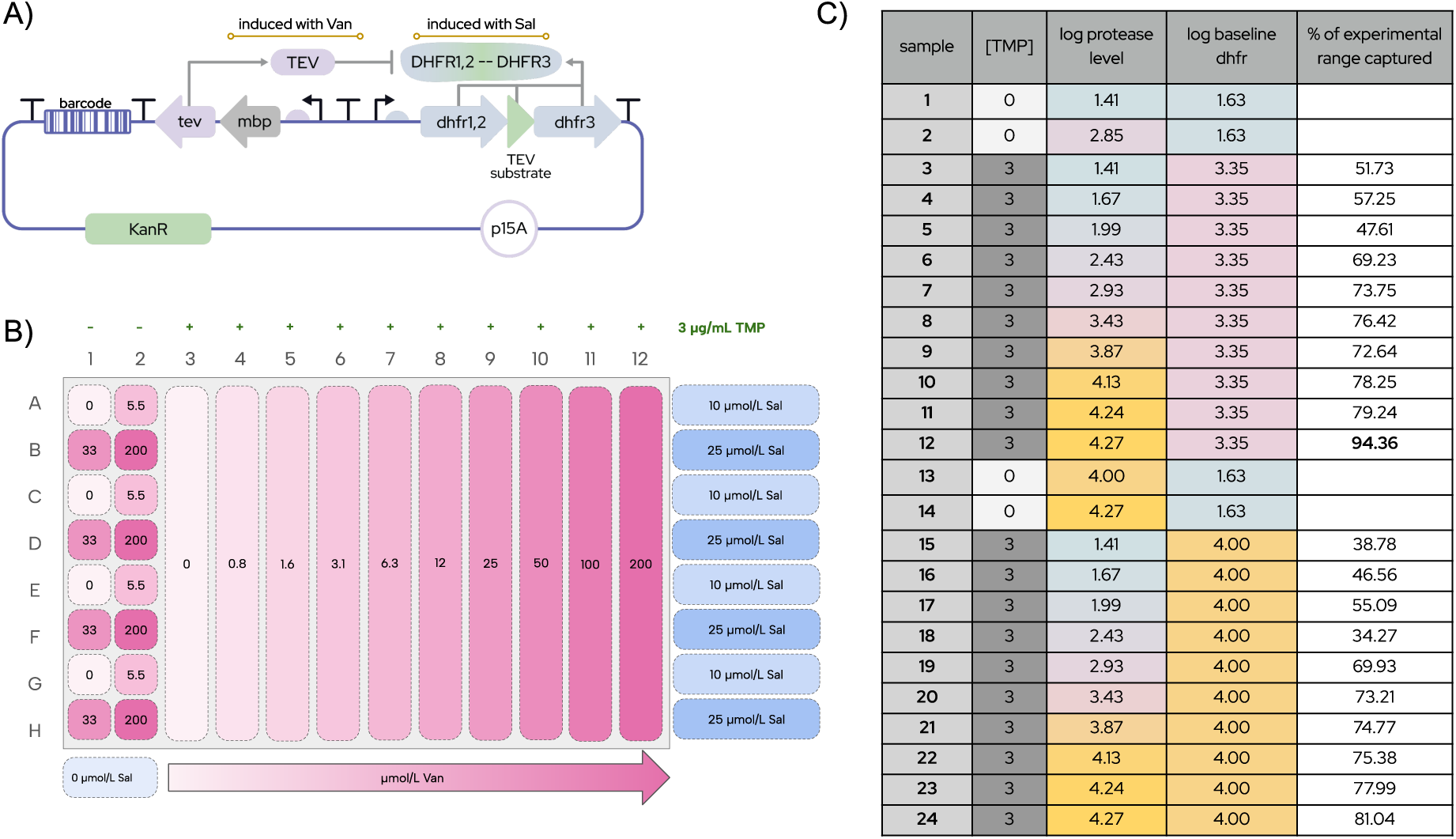
TEV Protease Experimental Setup. (**A**) schematic of plasmid circuit (Van= vanillic acid, Sal=salicylic acid, DHFR= dihydrofolate reductase, mbp=maltose binding protein); (**B**)Schematic of plate layout used in GROQ-seq experiment; (**C**) table of conditions tested and percentage of dynamic range captured (TMP=trimethoprim)

To systematically characterize TEV protease activity across a range of expression and selection regimes, we evaluated the SSVL and epPCR libraries under multiple experimental conditions. The TEV protease SSVL and epPCR libraries were evaluated across 24 conditions, each performed with four replicates on the same plate (Supplemental Figure 5B). Each condition varied along three parameters: TMP concentration, TEV protease expression level (vanillic acid concentration), and DHFR expression level (salicylic acid concentration). For each condition, the percent of dynamic range captured was calculated (Supplemental Figure 45). Based on this analysis, the figures in this Report focus on condition S12 which exhibited the highest dynamic range for variants with near-wild-type activity.

The dynamic range of the TEV protease assay was approximately 2 orders of magnitude. The TEV protease structure used for Figure 12B was from PDB entry 1LVM.

**For more details on the experiment, please refer to the Technical Bulletin for GROQ-seq TEV Protease Function Assay for details**^4^

## Notes

### Competing Interest Statement

The authors have declared no competing interest.

### Summary of Updates

I added an author that I accidentally forgot to include on the original submission (Douglas Densmore)

